# Patronus is the elusive plant securin, preventing chromosome separation by antagonizing separase

**DOI:** 10.1101/606285

**Authors:** Laurence Cromer, Sylvie Jolivet, Dipesh Kumar Singh, Floriane Berthier, Nancy De Winne, Geert De Jaeger, Shinichiro Komaki, Maria Ada Prusicki, Arp Schnittger, Raphael Guérois, Raphael Mercier

**Affiliations:** Institut Jean-Pierre Bourgin, UMR1318 INRA-AgroParisTech, Université Paris-Saclay, RD10, 78000 Versailles, France; Department of Plant Biotechnology and Bioinformatics, Ghent University, Ghent, Belgium, VIB Center for Plant Systems Biology, Ghent, Belgium; Department of Developmental Biology, Institute for Plant Sciences and Microbiology, University of Hamburg, 22609 Hamburg, Germany; Nara Institute of Science and Technology, Graduate School of Biological Sciences, 8916-5 Takayama, Ikoma, Nara 630-0192, Japan; Institute for Integrative Biology of the Cell (I2BC), Commissariat à l’Energie Atomique et aux Energies Alternatives (CEA), Centre National de la Recherche Scientifique (CNRS), Université Paris-Sud, CEA-Saclay, Gif-sur-Yvette, France

## Abstract

Chromosome distribution at anaphase of mitosis and meiosis is triggered by separase, an evolutionarily conserved protease. Separase must be tightly regulated to prevent the untimely release of chromatid cohesion and disastrous chromosome distribution defects. Securin is the key inhibitor of separase in animals and fungi, but has not been identified in other eukaryotic lineages. Here, we identified PATRONUS1 and PATRONUS2 (PANS1 and PANS2) as the Arabidopsis homologues of securin. Disruption of *PANS1* is known to lead to the premature separation of chromosomes at meiosis, and the simultaneous disruption of *PANS1* and *PANS2* is lethal. Here, we show that PANS1 targeting by the anaphase-promoting-complex is required to trigger chromosome separation, mirroring the regulation of securin. We showed that PANS1 acts independently from Shugosins. In a genetic screen for *pans1* suppressors, we identified *SEPARASE* mutants, showing that PANS1 and SEPARASE have antagonistic functions *in vivo*. Finally, we showed that the PANS1 and PANS2 proteins interact directly with SEPARASE. Altogether, our results show that PANS1 and PANS2 act as a plant securin. Remote sequence similarity was identified between the plant patronus family and animal securins, suggesting that they indeed derive from a common ancestor. Identification of patronus as the elusive plant securin illustrates the extreme sequence divergence of this central regulator of mitosis and meiosis.

## Introduction

Balanced segregation of chromosomes at both mitosis and meiosis requires that chromatid cohesion complexes be removed after proper chromosome alignment in the spindle. Inaccuracies in this process cause chromosome mis-segregation and aneuploidy, contributing to cancer and birth defects. Separase, which is conserved in fungi, animals and plants, triggers cohesion release by cleaving the kleisin subunit of the cohesin complex, opening the cohesin ring and allowing chromosome segregation (1). Securin is the primary regulator of separase activity in animals and fungi, forming a complex with separase and blocking substrate access to its active site. Securin, which is defined by its functional and biochemical properties, is a largely unstructured protein whose sequence is conserved only between closely related species (2–6). At the onset of anaphase, the anaphase-promoting-complex (also known as the cyclosome) (APC/C) triggers the degradation of securin, releasing separase activity and allowing chromosome separation. In vertebrates, the Cdk1-CyclinB1 complex is an additional regulator of separase activity (7). At meiosis I, the cleavage of cohesins is also spatially controlled: peri-centromeric cohesins are protected from separase cleavage by shugoshin-PP2A through dephosphorylation of the kleisin subunit (8, 9).

In Arabidopsis, separase is essential for cohesion release and chromosome segregation at both meiosis and mitosis (10, 11), APC/C regulates the progression of division (12, 13) and the shugoshins SGO1 and SGO2 and PP2A are required for cohesion protection at meiosis (14–16). However, securin is missing from the picture, raising the intriguing possibility that securin has been lost in the green lineage and that plant separase is regulated by a different mechanism.

We previously characterised the two paralogues *PATRONUS1* and *PATRONUS2* in Arabidopsis (PANS1, PANS2) (14). PANS1 is essential for the protection of sister chromatid cohesion between the two meiotic divisions, through an unknown mechanism. In the *pans1* mutant, sister chromatid cohesion is lost before metaphase II, leading to chromosome segregation defects at meiosis II (14, 17, 18). In addition, the *pans1* mutant also has a slight growth defect, which is exacerbated under stress conditions, associated with a certain level of mitotic defects and aneuploidy in somatic cells (14, 17, 19). The *pans2* mutant is indistinguishable from the wild type, but has synthetic lethal interaction with *pans1*, suggesting that PANS1 and PANS2 have an essential but redundant role at mitosis (14). PANS1 and PANS2 share 42% identity and encode proteins of unknown function. PANS1 has been shown to interact with APC/C through its D-box and KEN-box domains. Patronus proteins are well-conserved in dicots, one of the two major clades of flowering plants, with most species having one or two homologues (14). In monocots, the other major clade of flowering plants, PANS proteins share limited similarity with RSS1 (RICE SALT SENSITIVE 1, also known as Os02g39390), a protein that regulates cell cycle under stress conditions in rice (20). Although both PANS1, PANS2 and RSS1 clearly play important role in cell division, the molecular function of PANS1, PANS2 and RSS1 has remained elusive.

## Results

### Expression of destruction-box-less PATRONUS1 prevents cohesion release and chromosome segregation

The sequences of PANS proteins contain a conserved destruction-box (D-box) domain (RxxLxxxN), recognised by the anaphase-promoting complex (APC/C), which triggers the destruction of the targeted protein by the proteasome. A mutant version of PANS1 mutated in its D-box (PANS1ΔD; RxxL ➔ LxxV) loses its capacity to interact with the APC/C activator subunit CDC20 in yeast two-hybrid (Y2H) assays (14). Expression of *PANS1ΔD* under its own promoter is lethal, suggesting that PANS1 accumulation prevents plant development (14). To assess the function of the PANS1 D-box at meiosis, we expressed *PANS1ΔD* under the meiosis-specific promoter *DMC1*. Wild-type *PANS1* expressed under the *DMC1* promoter was able to complement the meiotic defects of the *pans1* mutant (n=2/4) (Figure S1). In contrast *pDMC1::PANS1ΔD* caused full sterility when transformed in wild-type or *pans1* plants (n=15/15 and 2/2, respectively). In addition, *pDMC1::PANS1ΔD* plants showed a variable growth defect, from barely developing plants to wild-type-like plants (Figure S2), suggesting that the *pDMC1* promoter can drive variable expression in somatic tissues, but all plants showed complete sterility. In *pDMC1::PANS1ΔD* plants, meiotic chromosome spreads did not reveal any defect in prophase or early metaphase I (Figure 1). However, among the 119 post-prophase cells observed, none showed the configuration typical of anaphase I, metaphase II, anaphase II or telophase II, suggesting that the meiotic cells do not progress beyond metaphase I. We observed normal metaphase I with five bivalents aligned on the metaphase plate (compare Figure 1F with Figure 1A), but also metaphase I with five over-stretched bivalents (Figure 1G). We also observed configurations resembling metaphase II, with chromosomes distributed in at least two groups separated by a dense band of organelles. However, five chromatid pairs aligned on each metaphase plates in the wild type (Figure 1D), whereas five bivalents aligned on two metaphase II plates in *pDMC1::PANS1ΔD* (e.g. 3 bivalents on one plate, and two on the other) (Figure 1I). At telophase II, we observed five nuclei, presumably each containing a decondensed bivalent. This observation suggests that *pDMC1::PANS1ΔD* abolishes the separation of homologous chromosomes at meiosis I. Immunostaining revealed that the bivalents in *pDMC1::PANS1ΔD* at a metaphase I- or metaphase II-like configuration were entirely decorated with the cohesin REC8, suggesting that the inhibition of chromosome separation is due to an incapacity to remove cohesins (Figure 2). We then observed the effect on *pDMC1::PANS1ΔD* on male meiocytes with live imaging of cells expressing a RFP-tagged tubulin (RFP:TUB4) and a GFP-tagged REC8 (21) (Figures 3-5 and supplemental movies S1-3). In the wild type (Movie S1), we observed the progression of male meiosis from prophase, with tubulin surrounding the nucleus (Figure 3A), to metaphase I with the formation of the spindle and alignment of the chromosomes (Figure 3C), and to anaphase I with the disappearance of the REC8 signal and reorganisation of the spindle (Figure 3D, 3E). The first division, from the end of prophase (nuclear break down, figure 3B) to the onset of anaphase I, lasted 38 ± 5 min (mean ± standard deviation, n=29 cells). When analysing *pDMC1::PANS1ΔD* with live imaging (Movies S2 and S3), we observed cells progressing normally from prophase to metaphase I (Figure 4A-C). However, the length of metaphase I was very variable, from 40 min (Movie S2, Figure 4) to more than 4 h (Movie S3). When anaphase occurred, as observed regarding microtubule reorganisation (Figure 4D-E, Movie S2), the REC8 signal was still detected. Further, five REC8:GFP bodies, presumably representing the five bivalents, were still observed at late anaphase I (arrows in Figure 4E), and following interphase, a spindle polymerised around each of the five REC8:GFP bodies (Figure 4F, Movie S2). Thus, live imaging confirmed that the expression of a D-box-less *PANS1* at meiosis prevents the release of REC8 and the separation of chromosomes.

**Figure 1.**
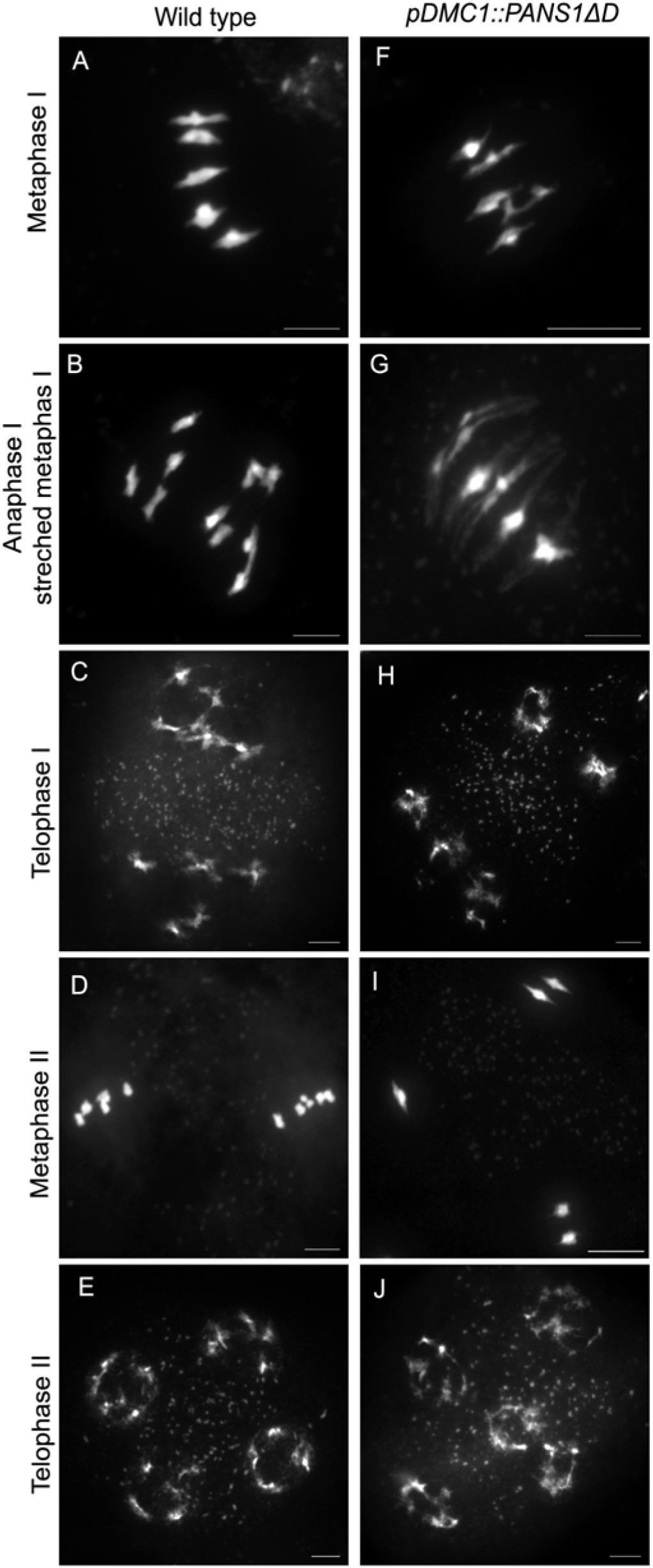
Meiotic spreads in the wild type (A-E) and *pDMC1::PANS1ΔD* (F-J). Male meiocytes were spread and stained with DAPI. Scale bar= 10 µM.

**Figure 2.**
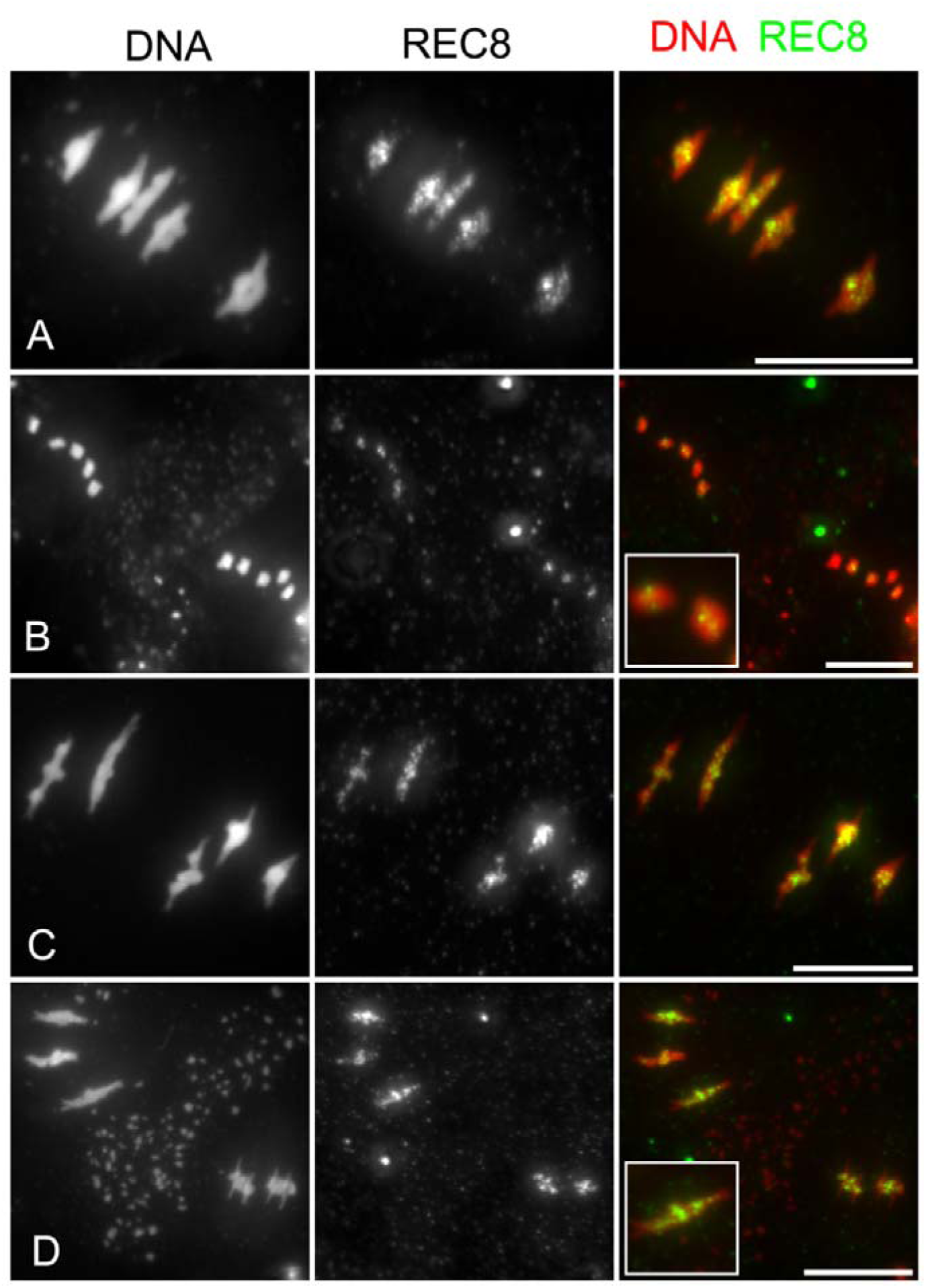
REC8 persists on chromosomes in *pDMC1::PANS1ΔD*. Immumocalisation of the cohesion REC8 was performed on meiotic chromosome spreads stained with DAPI (white). (A) Wild-type metaphase I. The bivalents under tension aligned on metaphase I are decorated with REC8. (B) Wild-type metaphase II. Five pairs of chromatids are aligned on both metaphase II plates. A faint REC8 signal is detected in the middle of each chromosome. (C) Metaphase I in *pDMC1::PANS1ΔD*. The bivalents are decorated with REC8, as in the wild type. (D) Aberrant metaphase II in *pDMC1::PANS1ΔD*. Two groups of bivalents are separated by an organelle band. The bivalents are under tension and are still decorated with REC8. Scale bar= 10 µM.

**Figure 3.**
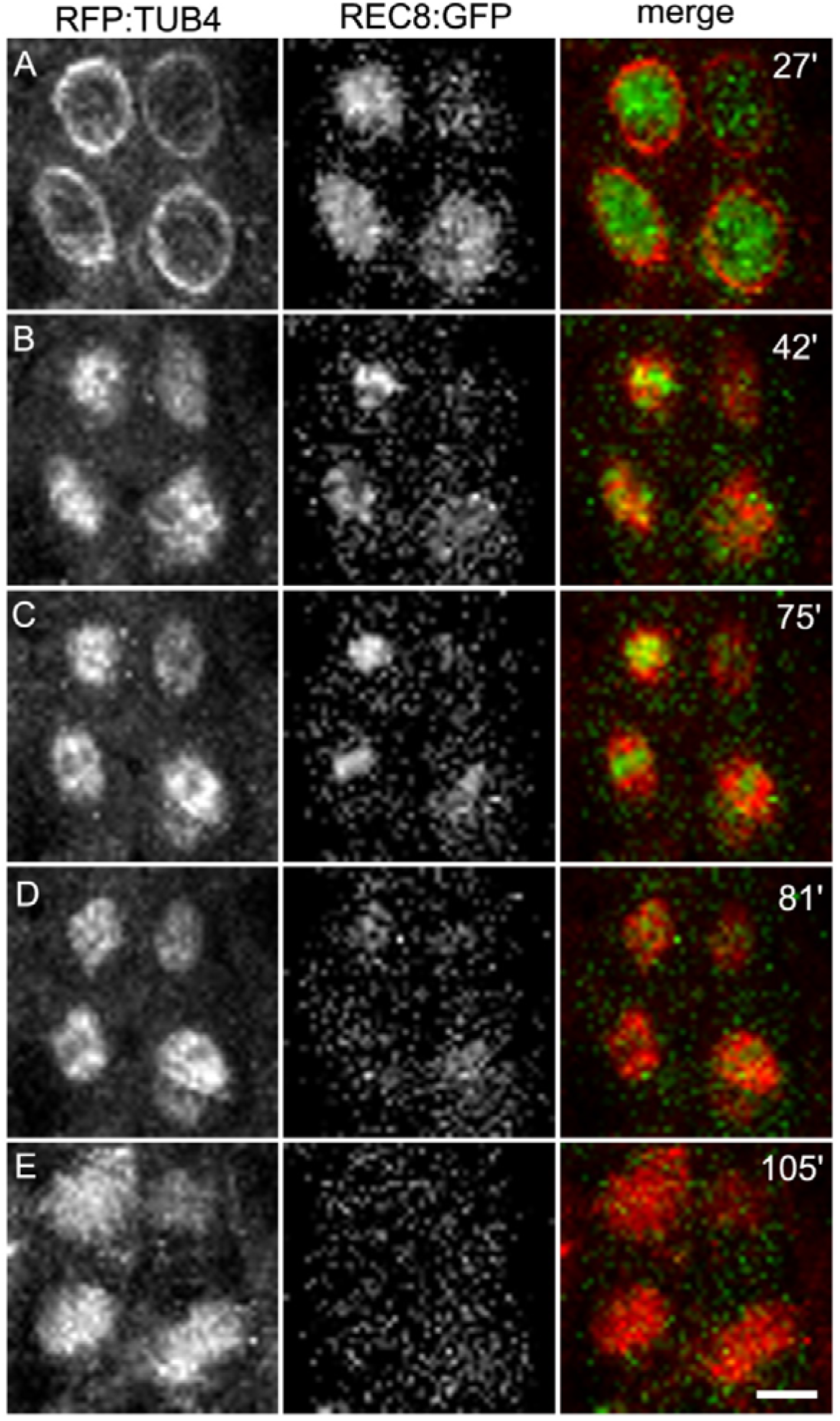
Time course of male meiosis I in wild type. Anthers at meiotic stages were observed in plant expressing the TUBULIN BETA CHAIN 4 fused with the RED FLUORESCENT PROTEIN (RFP:TUB4) and the cohesin REC8 fused with the GREEN FLUORESCENT protein (REC8:GFP). Images were acquired every 3 min (Movie S1). The time is indicated on the snapshots in minutes, starting with the beginning of the movie. (A) prophase. Tubulin surrounds the nucleus (B) at prometaphase. The nucleus disappears and the spindle is forming (C) Metaphase I chromosomes are aligned in the spindle. (D) Early anaphase I, the REC8:GPF signal suddenly decreases. (E) Late anaphase I. The spindle has reorganised. Scale bar= 10 µM.

**Figure 4.**
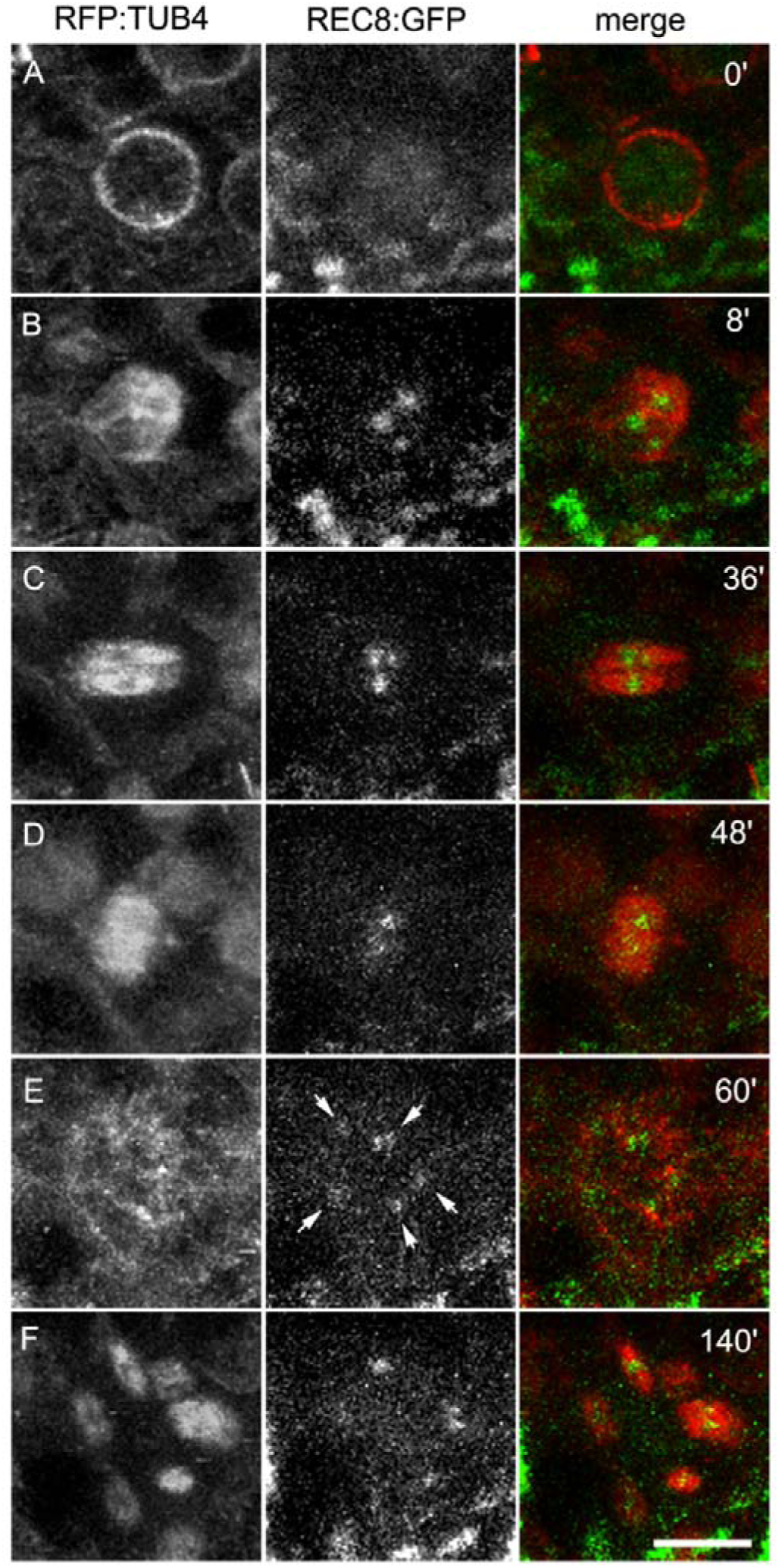
Time course of male meiosis in *pDMC1::PANS1ΔD*. Anthers at meiotic stages were observed in plants expressing RFP:TUB4, REC8:GFP and *pDMC1::PANS1ΔD*. Images were acquired every 4 min (Movie S2). The time is indicated on the snapshots in minutes, starting with the beginning of the movie (Movie S2). (A) Prophase. (B) Prometaphase. (C) Metaphase I). (D) Early anaphase I. (E) Abnormal late anaphase I with five REC8:GFP, indicated by arrows. (F) Aberrant metaphase II with five spindles. This time course corresponds to Movie S2. Movie S3 is an independent acquisition in the same background. Scale bar= 10 µM.

**Figure 5.**
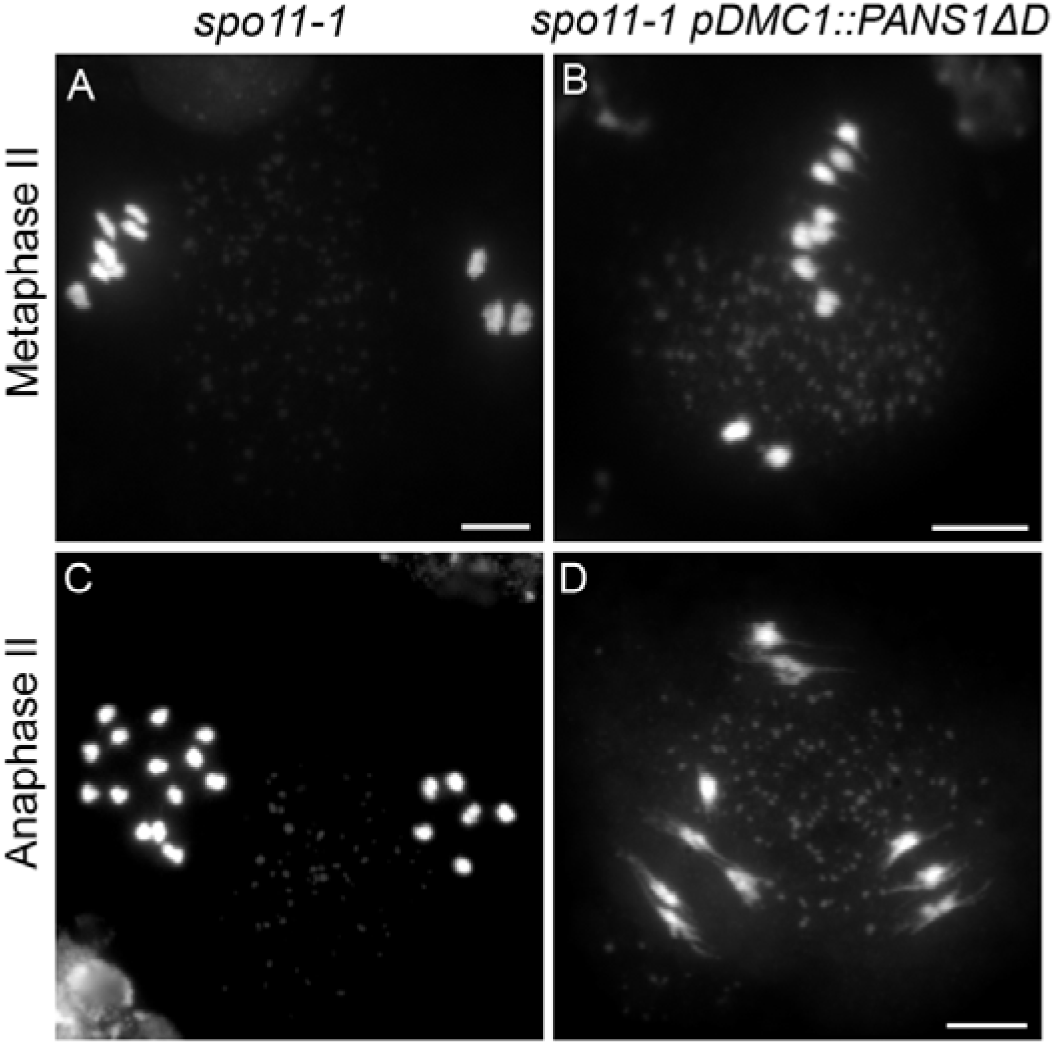
*pDMC1::PANS1ΔD* triggers arrest at metaphase II in *spo11-1*. Male meiocytes were spread and stained with DAPI. Scale bar= 10 µM. (A) Metaphase II in *spo11-1*. In this cell, three chromosomes are aligned on one plate, and seven on the other. (B) Metaphase II in *spo11-1* pDMC1::PANS1ΔD. Similarly, uneven numbers of chromosomes are aligned on the two metaphase plates. (C) Anaphase II in *spo11-1*. Anaphase II with separation of sister chromatids (3-3 and 7-7). (D) Pseudo-anaphase II in *spo11-1* pDMC1::PANS1ΔD with stretched chromosomes and no separation of sister chromatids. Scale bar= 5 µm.

In Arabidopsis *spo11-1* mutants, the formation of crossovers is eliminated and the resulting un-connected chromosomes (univalents) segregate randomly at anaphase I, and progress through metaphase II and anaphase II with segregation of sister chromatids (22) (Figure 5). Thus, the *spo11-1* mutation should allow segregation of chromosomes at meiosis I in *pDMC1::PANS1ΔD* and entry into meiosis II, making it possible to assess the effect of *PANS1ΔD* on chromatid pairs at anaphase II. Our results indicated that *spo11-1 pDMC1::PANS1ΔD* meiocytes progressed to metaphase II (Figure 5B), but appeared to be arrested at that stage with stretched chromosomes (Figure 5D), showing that *PANS1ΔD* can prevent chromosome segregation of both homologues at meiosis I and sister chromatids at meiosis II. The lethality of *PANS1ΔD* expressed under its own promoter suggests that it can also prevent chromosome segregation at mitosis. Altogether, these results suggest that targeting of PANS by APC/C is required for the release of cohesion and chromosome distribution at anaphase of mitosis and meiosis, mimicking the regulation and function of securin (7, 23, 24).

### PATRONUS1 acts independently of SHUGOSHINs

Shugoshin (SGO) is an evolutionarily conserved protein that protects sister chromatid cohesion against separase (8, 25). In Arabidopsis, both shugoshin genes (*SGO1* and *SGO2*) are involved in the protection of peri-centromeric cohesion during meiosis, with the corresponding double-mutant leading to the complete loss of sister chromatid cohesion at anaphase I (14), without apparent somatic function. The *pans1* mutant loses peri-centromeric cohesion during interkinesis. In both *pans1* and *sgo1 sgo2*, metaphase I appears normal with five aligned bivalents (14). To test if *PANS1* acts through or independently of *SGO*, we first combined *sgo1 sgo2* and *pans1* mutations (Figure 6 A-D). In the *pans1 sgo1 sgo2* triple mutant, only a small proportion of metaphase I showed five bivalents (11%, N=321 cells, Figure 6B), whereas the majority showed an almost complete (41%, Figure 6C) or complete (48%, Figure 6D) loss of cohesion, with 20 free chromatids. At diakinesis, the stage of prophase that immediately precedes metaphase I, five bivalents were systematically observed showing that crossovers occurred and sister chromatid cohesion was established (Figure 6A). Thus, in *pans1 sgo1 sgo2*, sister chromatid cohesion was lost at pro-metaphase I or early metaphase I, dismantling the bivalent into free chromatids. This result shows that PANS1 and SGOs act in parallel to protect sister chromatid cohesion at metaphase I. This also reveals that SGO1 and SGO2 protect cohesion not only at peri-centromeres, but also along the chromosome arms, redundantly with PANS1.

**Figure 6.**
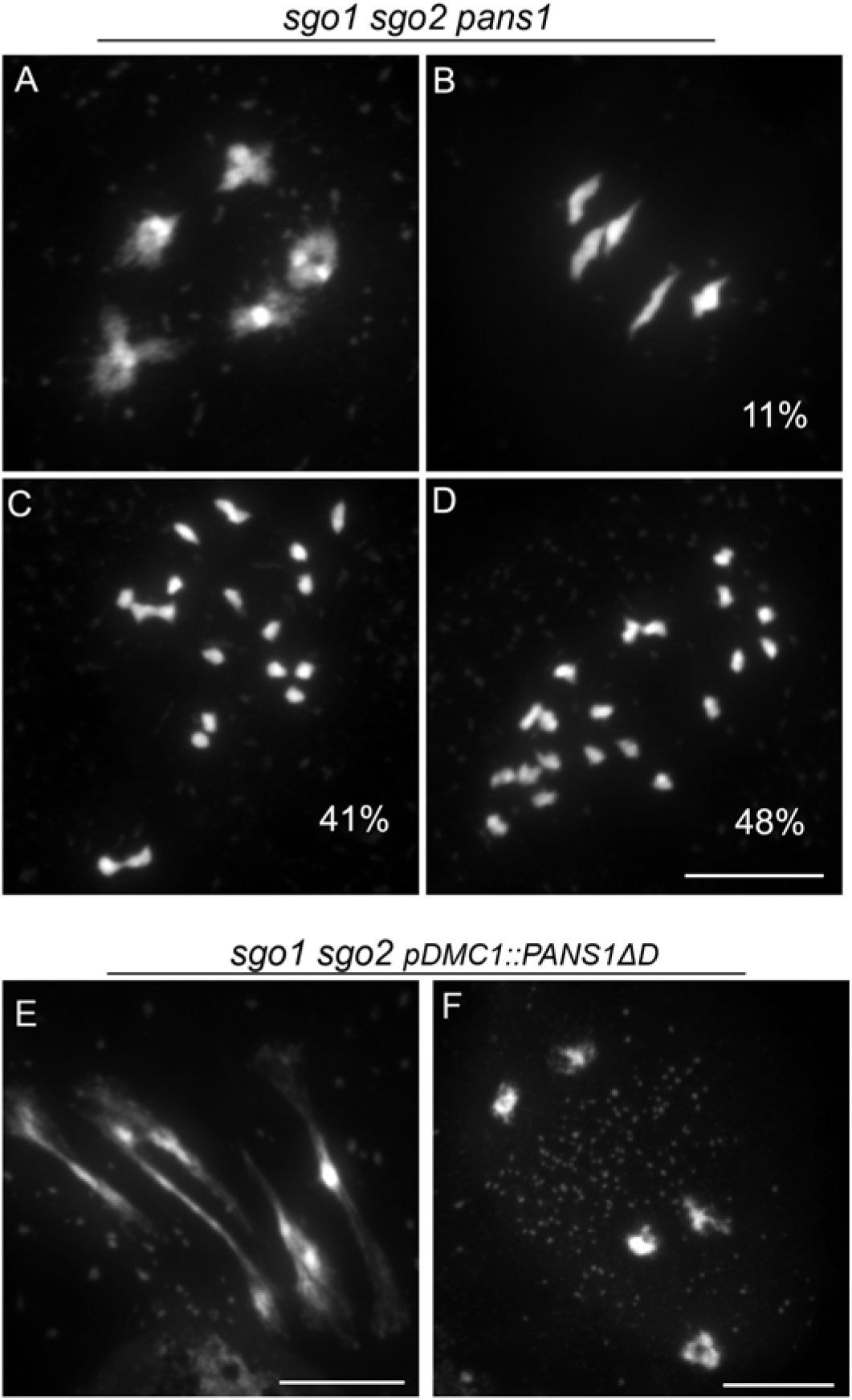
PANS1 acts independently of SGOs. (A-D) Chromosome spreads in *sgo1 sgo2 pans1* male meiosis. (A) Diakinesis. No defect is detected, five bivalents are observed. (B-D) Metaphases I, with five bivalents (B), almost complete loss of cohesion (C), or complete loss of cohesion leading to 20 free chromatids (D). (E-F) Chromosome spreads in *sgo1 sgo2 pDMC1::PANS1ΔD* male meiosis. (E) Metaphase I with stretched bivalents. (F) Aberrant metaphase II with bivalents. Scale bar = 10 µM

We then expressed *pDMC1::PANS1ΔD* in the *sgo1 sgo2* mutant (n=4 plants). In this context, meiotic chromosome spreads showed metaphase I with stretched bivalents (Figure 6E) and aberrant metaphase II with five bivalents (Figure 6F), similarly to what was observed when *pDMC1::PANS1ΔD* was expressed in the wild type (Figure 1). This shows that the expression of *PANS1ΔD* at meiosis prevents chromosome segregation even in the absence of SGOs, confirming that the PANS1 function is independent of SGOs.

### Mutations in *SEPARASE* can restore sister chromatid cohesion in *patronus1*

With the aim of identifying antagonists of *PATRONUS* and to shed light on its function, we set up a genetic suppressor screen. We took advantage of the root-growth defect of *pans1* when cultivated on medium supplemented with NaCl (14, 19). *pans1-1* seeds were mutated with ethyl methanesulfonate and the M2 families obtained by self-fertilisation of individual mutagenised plants were screened for (i) longer roots than *pans1-1* on NaCl medium and, subsequently, (ii) longer fruits than *pans1-1* after transfer to the greenhouse. Plants satisfying both criteria where identified in eight of the 200 independent families screened. Whole-genome sequencing of these plants revealed that four out of the eight suppressors had a missense mutation in the Arabidopsis *SEPARASE* gene (*AtESP;* At4g22970, Table S1), that we hereafter call *esp-S606N, esp-P1946L, esp-A2047T* and *esp-P2156S* (Figure 7-8, Figure S3). Bulk genome sequencing of a segregating population identified *esp-S606N* as the mutation most strongly linked to the growth phenotype among the mutations segregating in that line, further supporting the conclusion that the mutations in *ESP* are causal. The *esp-P1946L, esp-A2047T* and *esp-P2156S* mutations affected well-conserved residues of the protease domain of ESP (Figure S3). The S606 residue is in a less conserved helical domain and belongs to a stretch of serine, which may suggest regulation of ESP by phosphorylation. The previously described *esp-2* mutation is null and lethal (10), but can restore *pans1* growth in a dominant manner (Figure 7C), confirming that decreasing ESP activity can suppress the *pans1* somatic defect. Single-mutants *esp-S606N, esp-P1946L, esp-A2047T* and *esp-P2156S* are viable when homozygous, without apparent defects in growth or development, suggesting that they are hypomorphs. Quantification of root growth in *pans1* mutants segregating for the *esp-S606N* or *esp-P2156S* mutations showed that these mutations restore root growth in a semi-dominant and dominant manner, respectively (Figure 7D).

**Figure 7.**
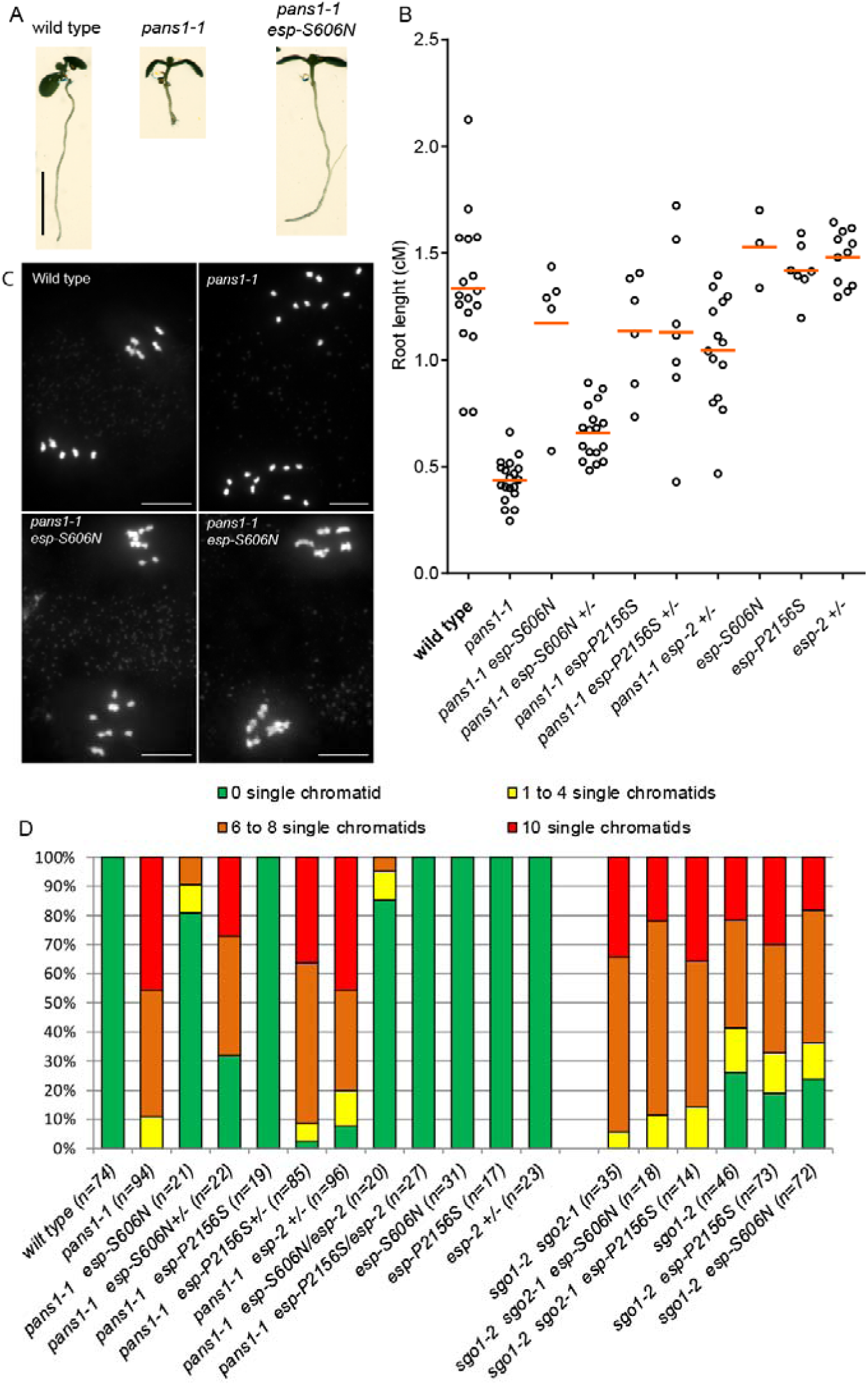
Mutations in *SEPARASE* suppress *pans1*. (A) Nine-day-old plantlets of wild type, *pans1-1* and *pans1-1 esp-S606N* grown on NaCl medium, as quantified in B. Scale bar =0.5 cM. (B) Quantification of roots length of nine-day-old plantlets. (C) Metaphase II plates of wild-type, *pans1-1* and *pans1-1 esp-S606N* plants as quantified in D. Scale bar =5 µM. (D) Quantification of sister chromatid cohesion. Metaphase II plates were sorted into classes according to the number of single chromatids detected, from 0 to 10. 0 means full cohesion as observed in wild type, 10 indicates a complete loss of cohesion. The number of metaphase plates analysed is indicated in parentheses.

**FIGURE 8.**
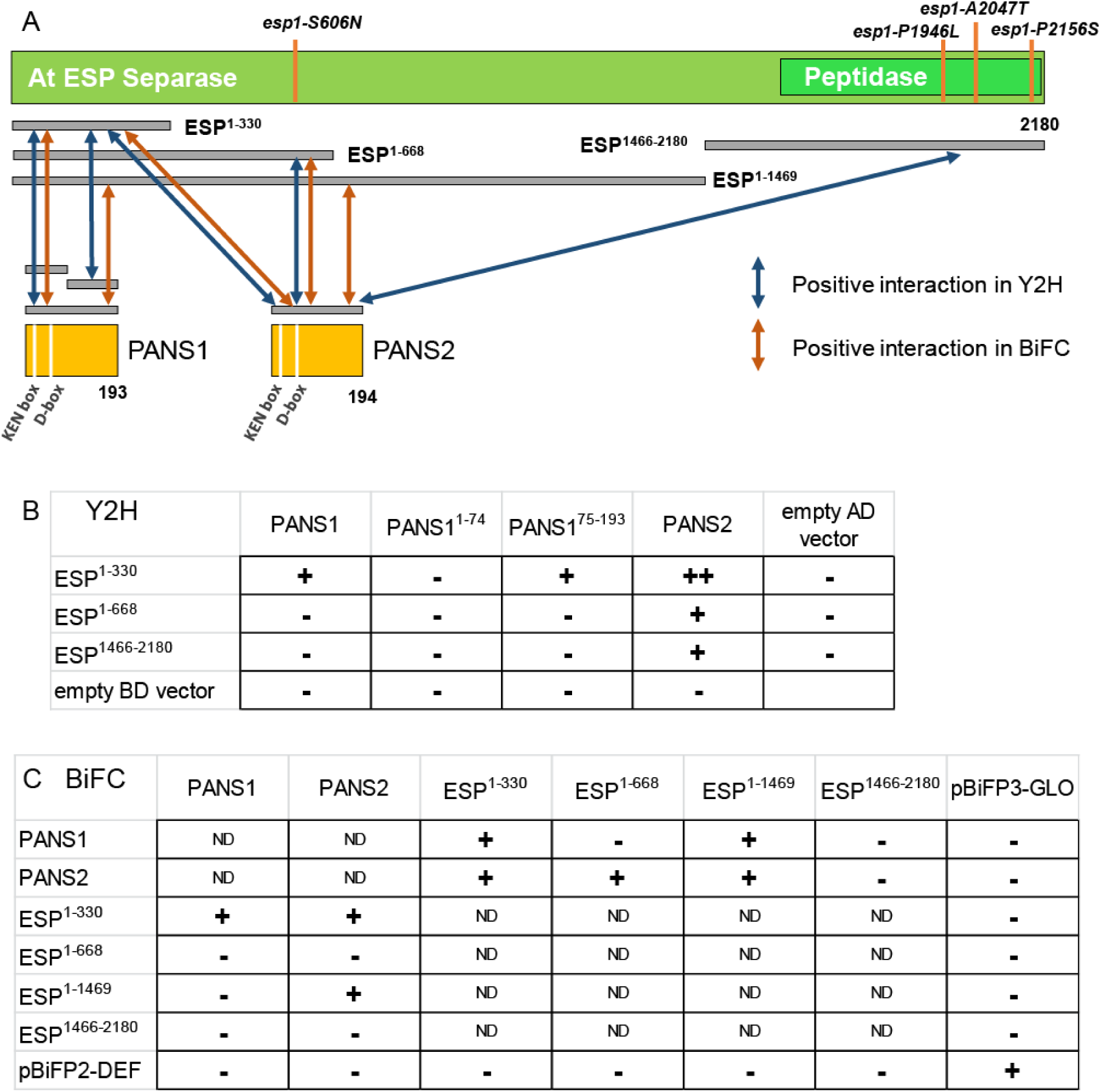
ESP and PANS1/2 interact in in yeast two-hybrid (Y2H) and BiFC assays. (A) Schematic representation of the ESP protein and its interactions with PANS1 and PANS2. The positions of the four mutations identified in this study and of the peptidase domain are indicated (see Figure S3). Positive interactions detected in Y2H and BiFC are indicated by arrows. (B) Results of Y2H essays. PANS1 and 2 proteins were fused with the activating domain (AD) and ESP was fused with the binding domain (BD). Growth on LWH is indicated by + and growth on LWHA is indicated by ++. Raw data are shown in Figure S4. (C) Results of BiFC essays. The C-terminus fusions with the C-YFP (pBiFP3) are listed in rows and the N-terminus fusion with N-YFP (pBiFP2) are listed in columns. + indicates the detection of a YFP signal (see Figure S5), - the absence of signal and ND, that the interaction was not tested.

To test if the mutations in *ESP* suppress the *pans 1* meiotic defects, we quantified sister chromatid cohesion at metaphase II (Figure 7C-D). Although cohesion is almost completely lost in *pans1* (Figure 7D, 10 free chromatids indicating complete absence of cohesion), it is partially restored in a semi-dominant manner by the *esp-S606N* mutation and fully restored by the *esp-P2156S* mutation in a recessive manner. The *esp-2* heterozygous mutation did not restore the *pans1* sister chromatid defect, but *pans1 esp-2*/*esp-P2156S* plants had restored cohesion, further confirming that *esp* mutations cause the suppression of the *pans1* phenotype. Thus, mutations in *ESP* can suppress the meiotic sister chromatid defect of *pans1*. In addition, *esp-S606N* and *esp-P2156S* mutations were not able to suppress the meiotic sister chromatid defect of *sgo1* or of *sgo1 sgo2* (Figure 7D). This result suggests that *PANS1* and *ESP* have a specific antagonistic function in regulating the release of sister chromatid cohesion at meiosis.

### PATRONUS interacts directly with SEPARASE and APC/C

To better understand the role of *PANS1*, we searched for interacting partners using pull-down protein purification coupled with mass spectrometry using over-expressed GS^rhino^-tagged PANS1 as bait in Arabidopsis cell culture (26). After filtering co-purified proteins for false positives (see Methods and (27)), we recovered peptides from PANS1 itself and a series of additional proteins in three replicate experiments (Table 1). We recovered 10 subunits of the APC/C complex, confirming previous findings (14). Most importantly, the PANS1 pull-down identified numerous peptides of SEPARASE (ESP1) in all three purification experiments, showing that PANS1 interacts with SEPARASE *in vivo*. Y2H assays and bimolecular fluorescent complementation (BiFC) experiments both confirmed that PANS1 and PANS2 interact with SEPARASE (Figure 8 and Figure S4). Y2H experiments with truncated PANS1 showed that the C-terminal half of PANS1 (which does not contain the conserved KEN and D boxes) is sufficient to mediate an interaction with SEPARASE. In yeast and animals, the securin C-terminal region also mediates the interaction with separase (4, 5) (Figure 9). The N-terminal domain of Arabidopsis SEPARASE showed the strongest interaction with PANS1 and PANS2. In yeast and human, securin interacts along the entire length of separase (4, 5, 7, 28).

**FIGURE 9.**
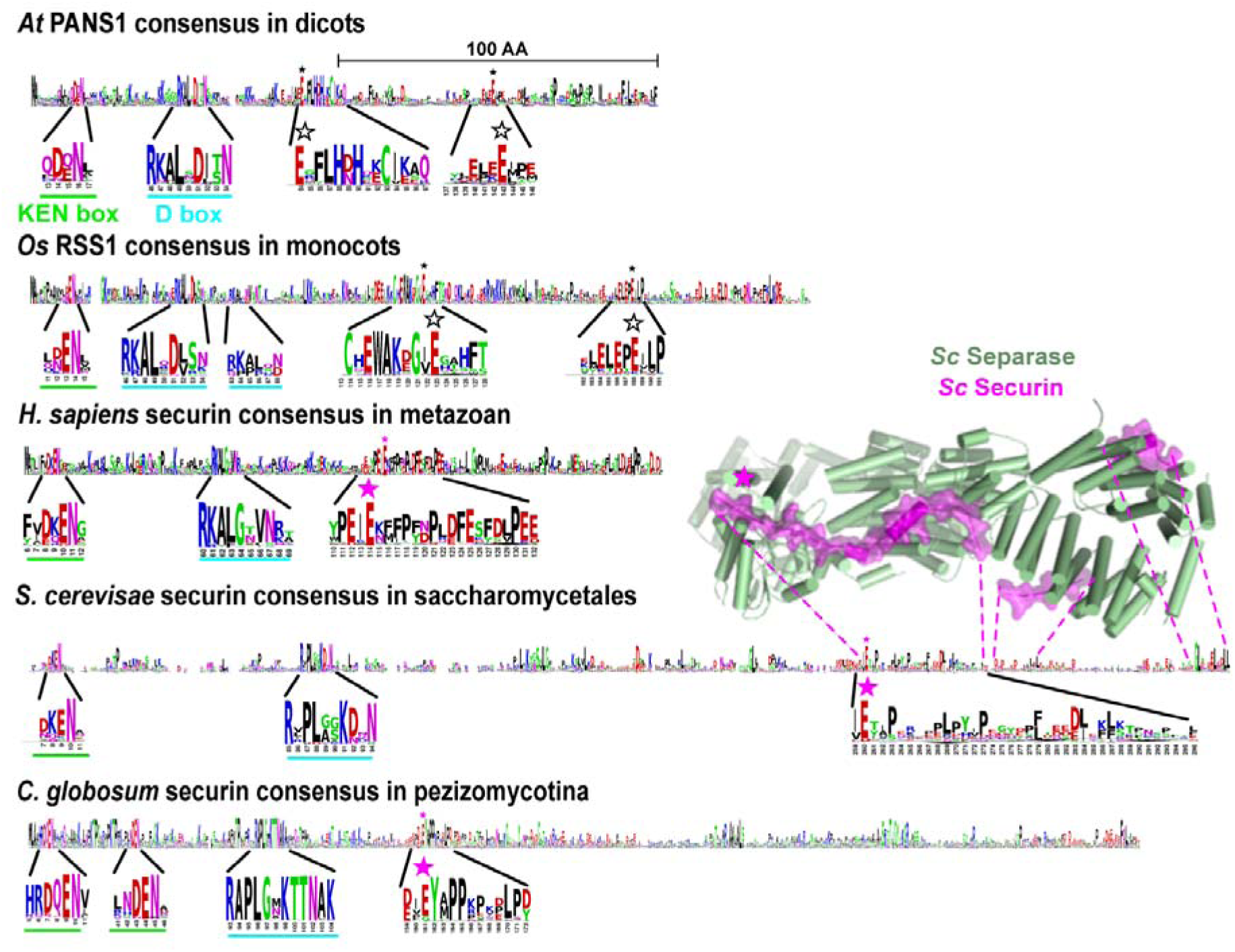
Conservation of securin organisation in eukaryotes. Sequence consensus of securin homologues obtained from the multiple sequence alignments of *Arabidopsis thaliana* PANS1 (PANS1_ARATH), *Oryza sativa* RSS1 (XP_015627424.1), *Homo sapiens* securin (AAC69752.1), *Saccharomyces cerevisiae* securin (NP_010398.3), *Chaetomium globosum* securin (XP_001225358.1) with their respective homologues represented using sequence logos (34). The homologues used to build the consensus span the dicot, monocot, metazoan, Saccharomycetale and Pezizomycotina clades for AtPANS1, OsRSS1, Hs, Sc and Cg securins, respectively. Sequence motifs corresponding to the KEN and D boxes are magnified and underlined in green and cyan, respectively. For all consensuses, the conserved stretches containing an invariant or almost-invariant glutamate residue (E) are magnified, with a star indicating the position of the conserved residue. Magenta stars report residues that were validated experimentally as acting as pseudo-substrates to inhibit separase activity (4, 5, 33). Unfilled black stars indicate the location of the potential pseudo-substrate glutamates in AtPANS1 and OsRSS1. The structure of Sc securin complexed to Sc separase is shown in magenta and green, respectively, with a magenta star indicating the position of the conserved glutamate and was obtained from the PDB code 5U1 T.

**Table 1.**

| Gene ID | function | Number of assigned peptide matches |  |  |  |
| --- | --- | --- | --- | --- | --- |
|  |  | IP1 | IP2 | IP3 | Total |
| AT3G14190 | PANS1 | 21 | 20 | 19 | 60 |
| AT4G22970 | SEPARASE | 27 | 31 | 36 | 94 |
| AT2G39090 | APC7 | 13 | 16 | 22 | 51 |
| AT5G05560 | APC1 | 13 | 16 | 20 | 49 |
| AT2G20000 | APC3b | 9 | 11 | 12 | 32 |
| AT3G48150 | APC8 | 9 | 11 | 10 | 30 |
| AT1G78770 | APC6 | 11 | 10 | 9 | 30 |
| AT4G21530 | APC4 | 3 | 6 | 12 | 21 |
| AT1G06590 | APC5 | 2 | 3 | 10 | 15 |
| AT2G04660 | APC2 | 3 | 2 | 3 | 8 |
| AT4G11920 | CCS52A2 | - | 3 | 2 | 5 |
| AT5G63135 | APC15 | - | - | 2 | 2 |
| AT1G48300 | Acyltransferase | 8 | 6 | 6 | 20 |
| AT3G13330 | PA200 | - | 2 | 3 | 5 |
| AT5G23900 | Ribosomal protein | 2 | 3 | - | 5 |
| AT3G07090 | Thiol peptidase | - | 2 | 2 | 4 |
| AT2G15270 | PRKRIP1 | 2 | - | - | 2 |

We previously showed that PANS1 interacts with the APC/C subunit CDC20 through its D box (14), and showed here that PANS2 also interacts with CDC20 (Figure S4). Yeast and animal securins have also been shown to bind directly CDC20 in a D-box-dependent manner (29, 30).

### PATRONUS has remote sequence similarity with securins

The experimental data presented above suggest that PATRONUS may be the elusive plant securin. We thus investigated the degree to which PATRONUS is conserved and if any sequence similarity can be detected between PATRONUS and animal or yeast securin.

Position-specific iterated BLAST (PSI-BLAST) against the green lineage identified homologues of PATRONUS and RSS1 in flowering plants, gymnosperms and basal vascular plants (e.g. mosses and ferns) as previously reported (14, 20). Further iterations then identified homologues in algae, including *Ostreococcus lucimarinus* (XP_001422400.1) (See Methods). Reciprocal PSI-BLAST analyses starting from the *O. lucimarinus* protein recovered the entire plant protein family, including RSS1, PATRONUS1 and PATRONUS2, reinforcing the conclusion that the identified proteins are homologues (Methods, Table S2). The slow convergence of the PSI-BLAST search likely arises from the high sequence divergence between the PATRONUS family members and from their intrinsically disordered character.

Next, we repeated the PSI-BLAST analyses using the *O. lucimarinus* sequence (XP_001422400.1) as bait and expanded the interrogation to the full eukaryotic tree. After eight iterations, a few proteins of unknown function from bivalves and gastropod species were identified, such as the bivalve mollusc *Mizuhopecten yessoensis* (XP_021355525.1). Remarkably, only a few PSI-BLAST iterations were sufficient to connect the latter sequence to homologues of securin in vertebrates, human and mouse securins. Other homologues experimentally probed as securin in *Caenorhabditis elegans*, *Drosophila melanogaster* or the fungi *Saccharomyces cerevisiae*, *Schizosaccharomyces pombe* and *Chaetomium thermophilum* were not retrieved in this search. Conversely, using these securins as bait of a PSI-BLAST search led to rapid convergence with homologues detected only in closely related species, suggesting that their sequences diverged too much to be recognised using this method. Hence, the fact that plant RSS1/PATRONUS can be connected to vertebrate securins in PSI-BLAST searches gives strong support to the suggestion that they share remote orthologous relationships, given that the lineage that led to plants has separated very early during eukaryotic evolution from the branch that led to animals and yeast (31).

Looking at the conservation patterns of the securin proteins within several clades, some striking features emerged (Figure 9 and Figure S4). In each clade (monocots, dicots, Saccharomycetales, Pezizomycotina, Metazoa), the same profile appeared: conserved KEN and D boxes in the N-terminal, and an invariable glutamic acid (E) within a relatively conserved patch. This glutamic acid has been shown to be pivotal for the inhibition of the separase cleavage site in the mammalian and the two fungus clades (pink stars in Figure 9), and is also shared with separase substrates (4, 7, 32, 33). In plants, two glutamic acids (E) are well conserved (Figure S4, unfilled stars in Figure 9) and are thus prime candidates important for separase inhibition. One is in the C-terminal third of the protein and is very well-conserved among the entire plant lineage (monocots, dicots, basal plants and algae). However, it is not conserved in some groups of species (e.g. the *Phalaenopsis* orchids, and several algae such as *Micractinium conductix*), making this candidate less likely. The other candidate glutamates lie in the middle of the proteins in domains conserved within dicots and within monocots in different regions (**E**-x-F-L-H-D/N-H and W–A–K/R–D/E/G–G–V/I–**E**, respectively; Figure 9 and S4), but of unknown function (14, 20). The monocot glutamate is extremely well-conserved in the Viridiplantae, ranging from monocots to very distant species such as algae (Figure S6), supporting its pivotal role. However, standard algorithms did not align the conserved E-containing patch from dicots with the conserved E-containing patch from monocots. We favour the scenario in which these E-containing domains represent the separase inhibition site, but that it has diverged too much in dicots to be properly aligned using the current algorithms.

## Discussion

Securin is the central regulator of chromosome distribution at both mitosis and meiosis in animals and fungi. Securin inhibits the separase protease, thus preventing untimely release of sister chromatid cohesion. At the onset of anaphase, APC/C degrades securin, releasing separase and thereby allowing chromosome distribution. However, to date, securin counterparts in plants have been elusive, suggesting that an alternative mechanism may be at play. We provide five pieces of strong evidence that patronus are securin homologues in plants. First, at meiosis, depletion of PANS1 leads to the premature release of cohesion (14); conversely, expression of an APC/C-insensitive PANS1 abolishes cohesion release and chromatid separation, mimicking the depletion of separase (10). If expressed constitutively, the APC/C-insensitive PANS1 is lethal. The single *pans1* mutation provokes some chromosome mis-segregation and aneuploidy (17, 19). No defects were detected in the *pans2* mutant, but the *pans1 pans2* double mutation is lethal (14). We propose that *PANS1* and *PANS1*, whose duplication is specific to Brassicaceae, have redundant functions, but that expression and/or activity levels of PANS1 are higher than those of PANS2. Our data support the conclusion that PATRONUS1 and/or PATRONUS 2 prevent cohesion release in both mitosis and meiosis and that degradation of PATRONUS by APC/C is required to lift this inhibition. Second, we showed that PANS1 controls chromosome segregation independently of SGOs, excluding the alternative hypothesis that patronus regulates shugoshin. Third, we demonstrated that the PATRONUS1 and 2 proteins interact directly with SEPARASE and APC/C. Fourth, using a forward genetic screen, we identified mutations in SEPARASE that suppress the defects of *patronus1* mutants, showing that PATRONUS1 and SEPARASE have antagonistic functions. Lastly, we identified remote sequence similarity between plant PATRONUS/RSS1 proteins and animal securins. Altogether, this strongly supports the conclusion that PATRONUS/RSS1 is the plant securin.

We propose that all plant PATRONUS homologues, including the rice RSS1 (35), represent plant securins. Consistent with this conclusion, RSS1 is expressed in dividing cells and is regulated by the APC/C in a D-box-dependent manner (20). Further, expression of RSS1 deleted of its N-terminal domain that contains D and KEN boxes is lethal. RSS1 functions in the regulation of the cell cycle, but the *rss1* mutant is viable and fertile (20). However, RSS1 has an uncharacterised paralogue in the rice genome (Os01g0898400) (20) (Figure S5), which may act redundantly with RSS1, as PANS1 and PANS2 do. RSS1 has been shown to interact with the PP1 phosphatase (20, 35, 36), suggesting that RSS1 is either regulated by PP1 or has an additional function than inhibiting securin.

It is intriguing that securin proteins have such a poorly conserved sequence although they play such a central role in cell division. The only conserved features are the presence of D and KEN boxes, which are involved in cell-cycle regulation, and a variable conserved patch containing glutamic acid (E), which is pivotal in separase inhibition. In the plant lineage, the phylogenetic analysis of PANS1/RSS1 suggests that, in addition to sequence divergence, the PANS family has a complex history of gene duplication/gene loss (14). Securin interacts with separase along all its intrinsically unstructured length (7) (Figure S9), and acts as a pseudo-substrate. It is likely that securin has no catalytic activity. One possibility is that the securin sequence simply drifts passively due to the absence of any selective pressure on its sequence, leading to relaxed purifying selection. Alternatively, securin being a key regulator of cell cycle, pivotal in development and the stress response, may evolve rapidly in response to selective pressures. In support of this idea, the rice *rss1* and the Arabidopsis *pans1* mutants are hypersensitive to a range of abiotic stresses (14, 19, 20) and intrinsically disordered protein regions are frequent targets of positive selection (37, 38). The rapid divergence of PATRONUS and its securin homologues may thus represent the accumulation of physiological and developmental responses to a constantly changing environment.

## Methods

### Plant materials and growth conditions

*Arabidopsis thaliana* plants were grown in greenhouses or growth chambers (16 h day/8 h night, 20°C, 70% humidity) and *Nicotiana benthamiana* in greenhouses (13 h day 25°C /11 h night 17°C). For *in vitro* culture, Arabidopsis seeds were surface sterilised for 10 min in 70% ethanol + 0.05% SDS and washed for 10 min in 70% ethanol and grown in petri dishes with culture medium (Gamborg B5 medium-Duchefa supplied with 0.16% BCP-bromocresol purple and 0.1% sucrose). The culture medium was supplemented with 80 mM NaCl for the root-growth experiment and with 100 µg/ml kanamycin sulfate (Euromedex) or 25 µg/ml hygromycin B (Duchefa) for selection of transformants. Arabidopsis transformation was performed using the floral dip method: *Agrobacterium tumefaciens* (EHA105 strain) was grown at 28–30°C to saturation, centrifuged and resuspended in a 1% sucrose 0.05% Silwet L-77 solution. This solution is used for dipping *A. thaliana* plants. Plants were maintained under glass overnight to increase humidity (39). The *A. tumefaciens* (C58C1 strain) was used for *Nicotiana benthamiana* leaf infiltration (40).

The *pans1-1* (Salk_070337), *Atsgo1-2* (SK2556), *Atsgo2-1* (line 34303), *spo11-1-3* (Salk_146172) and *esp-2* (Salk_037016) mutants were genotyped by PCR (30 cycles of 30 s at 94°C, 30 s at 56°C and 1 min at 72°C) and two primer pairs were used (Table S3). The first pair is specific to the wild-type allele and the second pair is specific to the left border of the inserted sequence. *pans1-1:* N570337U & N570337L, N570337L & LbSalk2. *Atsgo1-2*: SK2556U & SK2556L, SK2556U & pSKTail1. *Atsgo2-1*: SGO2U & SGO2L, SGO2U & GABI. *spo11-1-3*: N646172U & N646172L, N646172L & LbSalk2. *esp-2:* N537016U & N537016L, N537016U & LbSalk2.

The *esp-S606N* and *esp-P2156S* mutants were genotyped using PCR. The PCR products were digested by restriction enzymes. For *esp-S606N*, the 502 pb PCR product (esp606U & esp606L) was digested with the *Tas*I (Thermo Fisher Scientific) restriction enzyme at 65°C for 1 h. The wild-type allele yielded three DNA fragments (441 pb, 49 pb and 12 pb) and the *esp-S606N* allele yielded four DNA fragments (300 pb, 141 pb, 49 pb and 12 pb). For *esp-P2156S*, the 1040 pb PCR product (esp2156U & esp2156L) was digested with the *Hin*6I (Thermo Fisher Scientific) restriction enzyme at 37°C for 1 h. The wild-type allele yielded two DNA fragments (556 pb and 474 pb) and the *esp-P2156S* allele yielded one DNA fragment (1040 pb).

The RFP-tagged tubulin (RFP:TUB4) line and the GFP-tagged REC8 (REC8:GFP) were provided by AS. Homozygous lines for RFP:TUB4 and REC8:GFP were transformed using floral dip with the *pDMC1::PANS1ΔD* construct. Transformants were selected on petri dishes containing *in vitro* culture medium supplied with kanamycin.

### Suppressor screen

For the suppressor screen, homozygous *pans1-1* seeds were incubated for 17 h at room temperature in 5 mL of 0.3% (v/v) ethyl methanesulfonate (EMS) (SIGMA) with gentle agitation. EMS was neutralised by adding 5 mL 1M of sodium thiosulfate for 5 min. Then, 3 mL of water was added to make the seeds sink. The supernatant was removed and seeds were washed three times for 20 min with 15 mL of water. These M1 seeds were grown in the greenhouse and selfed to produce M2 seeds. M2 seeds were sterilised and sown in petri dishes containing *in vitro* culture medium supplemented with 80 mM NaCl (∼15 M2 per M1). M2 plants with long roots were transferred in the greenhouse and visually scored for fruit length. Mutants or mutant populations were sequenced using Illumina technology (GET platform https://get.genotoul.fr/). Mutations were identified using the MutDetect pipeline (41). The *esp-S606N* line identified in the suppressor screen was back-crossed with *pans1-1* and 52 plants were selected in the F2 population for having long roots on an NaCl medium. Bulk genome sequencing of these plants identified the *esp-S606N* mutation as the most strongly linked (36/41 mutant reads) to the phenotype among the mutation segregating in that line, supporting that it is causal.

### Meiotic chromosome spreads and immunolocalisation

Modified from (42). Inflorescences were harvested in 3:1 fixative (3 vol EtOH: 1 vol acetic acid). Fixative was replaced once. For slide preparation, inflorescences were washed twice in water and once in citrate buffer (10 mM tri-sodium-citrate, pH adjusted to 4.5 with HCl). They were digested for 3 h at 37°C in a moist chamber with a digestion mix (0.3% (w/v) pectolyase Y-23 (MP Biomedicals), 0.3% (w/v) Driselase™ (Sigma) 0.3% (w/v) cellulase (Onozuka R10) (Duchefa) 0.1% sodium azide in 10 mM citrate buffer). Three ∼0.5 mm washed buds were transferred on a slide in a drop of water and dilacerated with thin needles to generate a cell suspension. After adding 10 µl of 60% acetic acid, the slide was incubated on a hot block at 45°C for 1 min and the cell suspension was stirred with a hooked needle. Another 10 µl of 60% acetic acid was added and stirred during 1 more minute. The cell suspension drop was surrounded by fresh 3:1 fixative and the slide was rinsed with fixative. Dry slides were ready for DAPI staining and immunolocalisation.

For DAPI staining, a drop for DAPI solution (2 µg/mL in Citifluor™ AF1 (Agar scientific) was added. Slides were observed using a ZEISS Axio Imager Z2 microscope and Zen blue software. Images were acquired using a Plan-Apochromat 100x/1.40 Oil M27 objective, Optovar 1.25x Tubelens. DAPI was excited at λ 335-385 nm, and detected at λ between 420-470 nm.

For immunolocalisation, slides were microwaved in 10 mM citrate buffer pH 6 for 45 s at 850 W and immediately transferred to 0.1% Triton in PBS. Slides were incubated with primary antibodies diluted in 1% BSA in PBS for 48 h at 4 °C, then washed in 0.1% Triton in PBS three times for 15 min before adding the secondary antibodies in 1% BSA in PBS. After 1 h of incubation at 37 °C slides were washed in 0.1% Triton in PBS three times for 15 min and mounted in Vectashield antifade medium (Vector Laboratories) with 2 µg/ml DAPI. Slides were observed using a ZEISS Axio Imager Z2 microscope and Zen blue software. Images were acquired using a Plan-Apochromat 100x/1.40 Oil M27 objective, Optovar 1.25x Tubelens. DAPI was excited at λ 335-385 nm, and detected at λ between 420-470 nm. Green fluorescence was excited at λ 484-504 nm, and detected at λ between 517-537 nm. Red fluorescence was excited at λ 576-596 nm, and detected at λ between 612-644 nm. The rat anti-AtREC8 polyclonal antibody has been described by (14) and was used at a dilution of 1:250. The secondary antibody Alexa568 goat anti-rat (Thermo Fisher Scientific) was used at a dilution of 1:250.

### Live meiosis image acquisition and analysis

Live cell imaging was performed following Prusicki et al. (21) with modifications. Briefly, flowers buds of 0.4-0.6 mm were isolated on a slide. Buds were carefully dissected to isolate undamaged anthers. Anthers were transferred into a slide topped by a spacer (Invitrogen™ Molecular Probes™Secure-Seal™ Spacer, 8 wells, 9 mm diameter, 0.12 mm deep) filled with 10 µl of water, and covered by a coverslip. Time lapses were acquired using a Leica SP8 confocal laser-scanning and LAS X 3.5.0.18371 software.

Images were acquired using HC PL APO CS2 20x/0.75 IMM and HC PL APO CS2 63x/1.20 WATER objectives upon illumination of the sample with an argon laser and DPSS 561-nm. GFP was excited at λ 488 nm, and detected at λ between 494-547 nm. RFP was excited at λ 561 nm and detected at λ between 570-629 nm. Detection was performed using Leica HyD detectors. Time lapses were acquired as series of Z-stacks (between 15 to 20 μm distance). Interval time varied from 1.5 to 3 minutes depending on sample conditions. Deconvolution was performed using the lightning deconvolution option. Images processing were done with Fiji. Image drift was corrected by the Stack Reg plugin (Rigid Body option). Decrease in fluorescence was corrected using the bleach correction option (simple ratio- background intensity 0- or histogram matching).

### Plasmid construction

To generate the *pDMC1::PANS1* construct, the PANS1 genomic fragment was amplified using PCR with PANS1_XhoI and PANS1_SpeI primers. The amplification covered PANS1 from the ATG to 477pb after the stop codon. The PCR product was cloned by restriction digest with *Xho*I and *Spe*I into the pPF408 vector (43) to fuse the PANS1 genomic fragment with the *DMC1* promotor. The DMC1 promotor covered −2940 pb before ATG and +205 pb after ATG and has been amplified from the Landsberg eracta accession. The *pDMC1::PANS1* DNA fragment was amplified with DMC1_GTW and PANS1SpeI_GTW to introduce Gateway tail (Thermo Fisher Scientific) and cloned into pDONR207 to create pENTR-pDMC1::PANS1, on which directed mutagenesis was performed using the Stratagene Quick-change Site-Directed Mutagenesis Kit. The primer PANS1ΔD was used to create the pDMC1::PANS1ΔD. For plant transformation, the LR reaction was performed with the binary vector pGWB1(44).

For Y2H experiments, cDNA were amplified using the corresponding primers (see table S3). They were cloned using the Gateway cloning system (Thermo Fisher Scientific) into pDONR221 to generate pENTR clones and into pDEST22 (prey plasmid) and pDEST 32 (bait plasmid) Gateway®, ProQuest Thermo Fisher Scientific). To generate the pDONR221-ESP^1466–2180^ construct, a 4474 pb fragment of ESP cDNA was obtained by DNA synthesis (GeneArt-Thermo Fisher Scientific)(sequence available in Table S3). For BiFC experiments, the pENTR clones described above for Y2H were cloned by LR recombination reaction into the pBiFP2 and pBiFP3 vectors (40).

### Interaction Assay

#### Yeast two-hybrid assay

DNA product was cloned using Gateway (Invitrogen) into the pDONR221 vector (Invitrogen) to create pENTR. LR reactions were carried out on the pDEST32 (bait) and pDEST22 (prey) vectors (Invitrogen). Plasmids encoding the bait (pDEST32) and prey (pDEST22) were transformed into the yeast strain AH109 and Y187 (Clontech) by the LiAc method following the protocol in the MATCHMAKER GAL4 Two Hybrid System 3 manual (Clontech). The TDM protein self-interaction was used as positive controls (45). Transformed yeast cells were selected on synthetic drop-out (SD) plates without Leu (SD-L) for bait or without Trp (SD-W) for prey. Interactions between proteins were assayed using the mating method. The resulting diploid cells were selected on synthetic drop-out medium lacking a combination of amino acids, driven by the auxotrophy genes carried by the cloning vectors. Protein interactions were assayed by growing diploid cells on SD-LWH and SD-LWHA.

#### Bimolecular fluorescence complementation assays (BIFC)

Protein interactions were tested *in planta* using BiFC assays (46) in leaf epidermal cells of *N. benthamiana* plants expressing a nuclear CFP fused to histone 2B (Martin et al., 2009). N-terminal fusions, using the pENTR clones described above for Y2H, with two YFP complementary regions (YFPN + YFPC) were co-infiltrated in *N. benthamiana* leaves and scored after 3 or 4 days for fluorescence as described in [41]. YFPN::DEF and As positive controls, we co-expressed YFPN-DEF and YFPC-GLO, two interacting components of the *Anthirrinum majus* MADS box transcription factors DEFICIENS and GLOBOSA (47). Each experiment was replicated at least twice, each corresponding to the infiltration of two different plants.

Observations were made using a Leica SP8 confocal laser-scanning microscope. Optical sections were collected with a Leica HCX PL APO CS2 20.0×0.70 IMM UV water objective upon illumination of the sample with a 514 nm argon laser line with an emission band of 520–560 nm for the YFP or with a 458 nm argon laser line with an emission band of 463–490 nm for the CFP. Detection were performed using Leica HyD detectors. The specificity of the YFP signal was systematically checked by determining the fluorescence emission spectrum between 525 and 600 nm with a 10 nm window and under an excitation at 514 nm. Images were processed using Leica LASX and Adobe Photoshop software.

### Pull-downs

Three pull-downs on Arabidopsis cell suspension culture expressing N-terminally GS^rhino^ tagged PANS1 were performed as described (Van Leene et al., 2019, Nature Plants, in press). On-bead digested samples were analysed on a Q Exactive mass spectrometer (Thermo Fisher Scientific) and co-purified proteins were identified using standard procedures (27). After identification, the protein list was filtered versus a list of non-specific proteins, assembled similarly as described (27). True interactors that might have been filtered out due to their presence in the list of non-specific proteins were selected by means of semi-quantitative analysis using the average normalised spectral abundance factors (NSAF) of the identified proteins in the PANS1 pull-downs. Proteins identified with at least two peptides in at least two experiments, showing high (at least 10-fold) and significant [-log_10_(*p*-value(T-test)) ≥10] enrichment compared to calculated average NSAF values from a large dataset of pull downs with non-related bait proteins, were selected.

### Protein sequence analyses

Searches for homologues of AtPANS1 (PANS1_ARATH) were performed using PSI-BLAST(48) against the nr database using e-values thresholds from 1E-3 to 1E-5. Multiple sequence alignments were calculated using MAFFT (49) and represented using Jalview (50). PSI-BLAST searches starting from AtPANS1 established the homologous relationship with OsRSS1 (XP_015627424.1) after four iterations, but further iterations were not successful in identifying plant homologues in clades other than monocots and dicots. Repeating the search from OsRSS1 on plant sequences excluding dicots helped identify homologues in mosses. These sequences were used in turn as inputs to search for the most likely homologues of AtPANS1 in green algae and one sequence in *O. lucimarinus* was detected (XP_001422400.1) as a potential candidate. Reciprocal PSI-Blast analyses starting from the *O. lucimarinus* protein against the nr sequence database restricted to the *Viridiplantae* clade recovered OsRSS1 as a significant match after seven iterations ad the Arabidopsis PATRONUS1 and 2 after 13 iterations.

PSI-BLAST analyses using (XP_001422400.1) also detected as significant homologues several sequences from bivalves such as *M. yessoensis* (XP_021355525.1) after 10 iterations without apparent divergence of the sequence profile. PSI-Blast from *M. yessoensis* (XP_021355525.1) against metazoan sequences integrated homologues of securin in vertebrates after four iterations including those of *Homo sapiens*, establishing a transitivity link from Arabidopsis PANS1 to human securin passing through green algae and bivalve homologues. The sequence accession numbers retrieved after the 10 iterations of PSI-BLAST from the *O. lucimarinus* sequence are listed in Table S2.

## Acknowledgements

We are grateful to Mathilde Grelon, Christine Mezard and Eric Jenczewski for critical reading of the manuscript. The Institute Jean-Pierre Bourgin benefits from the support of the LabEx Saclay Plant Sciences-SPS (ANR-10-LABX-0040-SPS). This work was financially supported by a European Research Council Grant ERC 2011 StG 281659 (MeioSight) to R.M., the Fondation Simone et Cino Del Duca to R.M. and the CEFIPRA project SMOKY to R.M.. Furthermore, this work was supported by the European Union Marie-Curie “COMREC” network FP7 ITN-606956 to M.A.P. and A.S. In addition, core funding from the University of Hamburg to A.S. is gratefully acknowledged.

## References

1. Kamenz J, Hauf S (2016) Time To Split Up: Dynamics of Chromosome Separation. Trends Cell Biol 27(1):42–54.

2. Hornig NCD, Knowles PP, McDonald NQ, Uhlmann F (2002) The dual mechanism of separase regulation by securin. Curr Biol 12(12):973–82.

3. Hauf S, Waizenegger IC, Peters JM (2001) Cohesin cleavage by separase required for anaphase and cytokinesis in human cells. Science (80-) 293(5533):1320–3.

4. Luo S, Tong L (2017) Molecular mechanism for the regulation of yeast separase by securin. Nature 542(7640):255–259.

5. Boland A, et al. (2017) Cryo-EM structure of a metazoan separase-securin complex at near-atomic resolution. Nat Struct Mol Biol 24(4):414–418.

6. Zou H, McGarry TJ, Bernal T, Kirschner MW (1999) Identification of a vertebrate sister-chromatid separation inhibitor involved in transformation and tumorigenesis. Science (80-) 285(5426):418–421.

7. Luo S, Tong L (2018) Structural biology of the separase–securin complex with crucial roles in chromosome segregation. Curr Opin Struct Biol 49:114–122.

8. Kitajima TS, Kawashima S a, Watanabe Y (2004) The conserved kinetochore protein shugoshin protects centromeric cohesion during meiosis. Nature 427(6974):510–7.

9. Miyazaki S, Kim J, Sakuno T, Watanabe Y (2017) Hierarchical Regulation of Centromeric Cohesion Protection by Meikin and Shugoshin during Meiosis I. Cold Spring Harb Symp Quant Biol LXXXII:033811.

10. Liu Z, Makaroff CA (2006) Arabidopsis separase AESP is essential for embryo development and the release of cohesin during meiosis. Plant Cell 18(5):1213–25.

11. Moschou PN, Bozhkov PV. (2012) Separases: Biochemistry and function. Physiol Plant 145(1):67–76.

12. Fülöp K, et al. (2005) Arabidopsis anaphase-promoting complexes: Multiple activators and wide range of substrates might keep APC perpetually busy. Cell Cycle 4(8):1084–1092.

13. Kevei Z, et al. (2011) Conserved CDC20 cell cycle functions are carried out by two of the five isoforms in Arabidopsis thaliana. PLoS One 6(6):e20618.

14. Cromer L, et al. (2013) Centromeric cohesion is protected twice at meiosis, by SHUGOSHINs at anaphase i and by PATRONUS at interkinesis. Curr Biol 23(21):2090–2099.

15. Zamariola L, et al. (2013) SGO1 but not SGO2 is required for maintenance of centromere cohesion in Arabidopsis thaliana meiosis. Plant Reprod 26(3):197–208.

16. Yuan G, et al. (2018) PROTEIN PHOSHATASE 2A B’α and β Maintain Centromeric Sister Chromatid Cohesion during Meiosis in Arabidopsis. Plant Physiol 178(1):317–328.

17. Zamariola L, et al. (2014) SHUGOSHINs and PATRONUS protect meiotic centromere cohesion in Arabidopsis thaliana. Plant J 77(5):782–94.

18. Singh DK, Spillane C, Siddiqi I (2015) PATRONUS1 is expressed in meiotic prophase I to regulate centromeric cohesion in Arabidopsis and shows synthetic lethality with OSD1. BMC Plant Biol 15(1):1–11.

19. Juraniec M, et al. (2016) Arabidopsis COPPER MODIFIED RESISTANCE1/PATRONUS1 is essential for growth adaptation to stress and required for mitotic onset control. New Phytol 209(1):177–191.

20. Ogawa D, et al. (2011) RSS1 regulates the cell cycle and maintains meristematic activity under stress conditions in rice. Nat Commun 2:278.

21. Prusicki MA, et al. (2018) Live cell imaging of meiosis in Arabidopsis thaliana - a landmark system. bioRxiv:446922.

22. Grelon M, Vezon D, Gendrot G, Pelletier G (2001) AtSPO11-1 is necessary for efficient meiotic recombination in plants. EMBO J 20(3):589–600.

23. Cohen-Fix O, Peters JM, Kirschner MW, Koshland DE (1996) Anaphase initiation in saccharomyces cerevisiae is controlled by the APC-dependent degradation of the anaphase inhibitor Pds1p. Genes Dev 10(24):3081–3093.

24. Funabiki H, et al. (1996) Cut2 proteolysis required for sister-chromatid separation in fission yeast. Nature 381(6581):438–441.

25. Clift D, Marston AL (2011) The role of shugoshin in meiotic chromosome segregation. Cytogenet Genome Res 133(2–4):234–42.

26. Bontinck M, et al. (2018) Recent Trends in Plant Protein Complex Analysis in a Developmental Context. Front Plant Sci 9(May):1–14.

27. Van Leene J, et al. (2015) An improved toolbox to unravel the plant cellular machinery by tandem affinity purification of Arabidopsis protein complexes. Nat Protoc 10(1):169–87.

28. Viadiu H, Stemmann O, Kirschner MW, Walz T (2005) Domain structure of separase and its binding to securin as determined by EM. Nat Struct Mol Biol 12(6):552–553.

29. Hilioti Z, Chung YS, Mochizuki Y, Hardy CFJ, Cohen-Fix O (2001) The anaphase inhibitor Pds1 binds to the APC/C-associated protein Cdc20 in a destruction box-dependent manner. Curr Biol 11(17):1347–52.

30. Kitagawa R, Law E, Tang L, Rose AM (2002) The Cdc20 homolog, FZY-1, and its interacting protein, IFY-1, are required for proper chromosome segregation in Caenorhabditis elegans. Curr Biol 12(24):2118–2123.

31. Harashima H, Dissmeyer N, Schnittger A (2013) Cell cycle control across the eukaryotic kingdom. Trends Cell Biol 23(7):345–356.

32. Nagao K, Yanagida M (2006) Securin can have a separase cleavage site by substitution mutations in the domain required for stabilization and inhibition of separase. Genes to Cells 11(3):247–260.

33. Lin Z, Luo X, Yu H (2016) Structural basis of cohesin cleavage by separase. Nature 532(7597):131–134.

34. Crooks GE, Hon G, Chandonia J-M, Brenner SE (2004) WebLogo: a sequence logo generator. Genome Res 14(6):1188–90.

35. Ogawa D, Morita H, Hattori T, Takeda S (2012) Molecular characterization of the rice protein RSS1 required for meristematic activity under stressful conditions. Plant Physiol Biochem 61:54–60.

36. Ebel C, Hanin M (2016) Maintenance of meristem activity under stress: Is there an interplay of RSS1-like proteins with the RBR pathway? Plant Biol 18(2):167–170.

37. Afanasyeva A, Bockwoldt M, Cooney CR, Heiland I, Gossmann TI (2018) Human long intrinsically disordered protein regions are frequent targets of positive selection. Genome Res 28(7):975–982.

38. Nilsson J, Grahn M, Wright APH (2011) Proteome-wide evidence for enhanced positive Darwinian selection within intrinsically disordered regions in proteins. Genome Biol 12(7):R65.

39. Clough SJ, Bent AF (1998) Floral dip: a simplified method for Agrobacterium-mediated transformation of Arabidopsis thaliana. Plant J 16(6):735–43.

40. Azimzadeh J, et al. (2008) Arabidopsis TONNEAU1 proteins are essential for preprophase band formation and interact with centrin. Plant Cell 20(8):2146–59.

41. Girard C, et al. (2014) FANCM-associated proteins MHF1 and MHF2, but not the other Fanconi anemia factors, limit meiotic crossovers. Nucleic Acids Res 42(14):9087–9095.

42. Ross KJ, Fransz P, Jones GH (1996) A light microscopic atlas of meiosis in Arabidopsis thaliana. Chromosom Res 4(7):507–16.

43. Siaud N, et al. (2004) Brca2 is involved in meiosis in Arabidopsis thaliana as suggested by its interaction with Dmc1. EMBO J 23(6):1392–401.

44. Nakagawa T, et al. (2007) Development of series of gateway binary vectors, pGWBs, for realizing efficient construction of fusion genes for plant transformation. J Biosci Bioeng 104(1):34–41.

45. Cifuentes M, et al. (2016) TDM1 Regulation Determines the Number of Meiotic Divisions. PLoS Genet 12(2):e1005856.

46. Hu C-D, Chinenov Y, Kerppola TK (2002) Visualization of interactions among bZIP and Rel family proteins in living cells using bimolecular fluorescence complementation. Mol Cell 9(4):789–98.

47. Schwarz-Sommer Z, et al. (1992) Characterization of the Antirrhinum floral homeotic MADS-box gene deficiens: evidence for DNA binding and autoregulation of its persistent expression throughout flower development. EMBO J 11(1):251–63.

48. Altschul SF, et al. (1997) Gapped BLAST and PSI-BLAST: a new generation of protein database search programs. Nucleic Acids Res 25(17):3389–402.

49. Katoh K, Standley DM (2013) MAFFT multiple sequence alignment software version 7: improvements in performance and usability. Mol Biol Evol 30(4):772–80.

50. Waterhouse AM, Procter JB, Martin DMA, Clamp M, Barton GJ (2009) Jalview Version 2--a multiple sequence alignment editor and analysis workbench. Bioinformatics 25(9):1189–91.

