## Supplementary material for "Patronus is the elusive plant securin, preventing chromosome separation by antagonizing separase": Figure S1

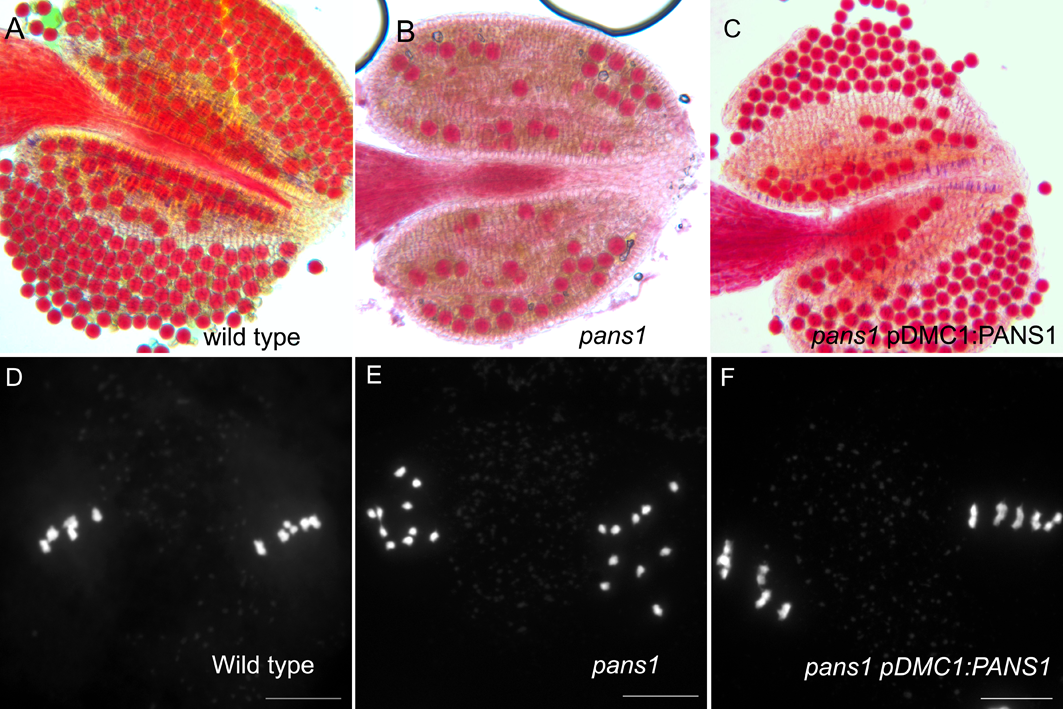

**Figure S1. pDMC1::PANS1 complements the *pans1* mutant**

(A-C) Alexander staining of anthers, with viable pollen appearing in red. In contrast to the wild type (A), a large proportion of pollen grains were dead in *pans1* anthers (B). *pans1* plants transformed by *pDMC1::PANS1* (C) have restored pollen viability (n=2/4). (D-F) Chromosome spreads at metaphase II. In the wild type (D), the two metaphase plates contain five pairs of chromatids. In *pans1* (E), the two metaphase plates contain ten free chromatids. In *pans1* transformed by pDMC1:PANS1 (F), chromatid cohesion is restored (n=2/4).
