## Supplementary material for "Patronus is the elusive plant securin, preventing chromosome separation by antagonizing separase": Figure S2

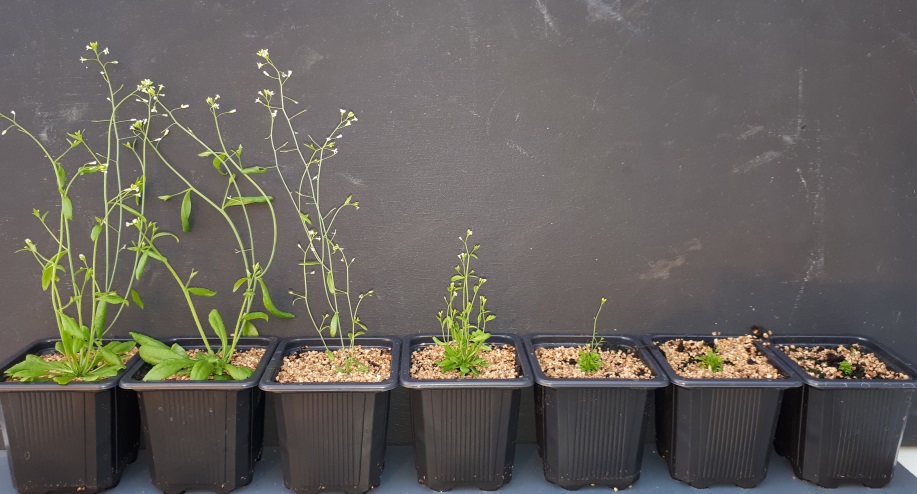

**FIGURE S2. *pDMC1::PANS1ΔD* have variable growth defects**

*pDMC1::PANS1ΔD* was transformed in wild type. These seven primary transformants were grown in parallel for five weeks, and show a range of growth defects.
