## Supplementary material for "Patronus is the elusive plant securin, preventing chromosome separation by antagonizing separase": Figure S3

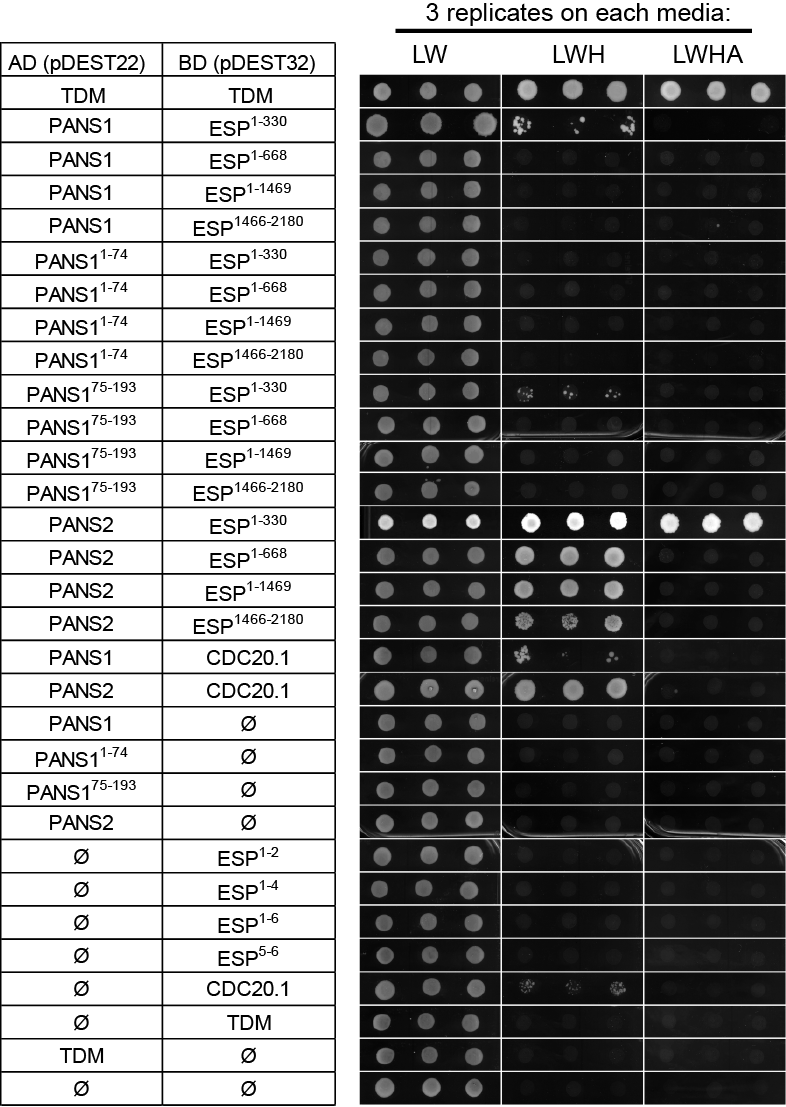

**Figure S4. Yeast two-hybrid assays**

Proteins of interest were fused with the GAL4 activating domain (AD) or the GAL4 binding domain (BD). TDM self-interaction was used as positive controls (Cifuentes et al., 2016). Protein interactions were assayed by growing diploid cells on SD-LW (control), SD-LWH (weak interaction) and SD-LWHA (strong interaction).
