## Supplementary material for "Patronus is the elusive plant securin, preventing chromosome separation by antagonizing separase": Figure S5

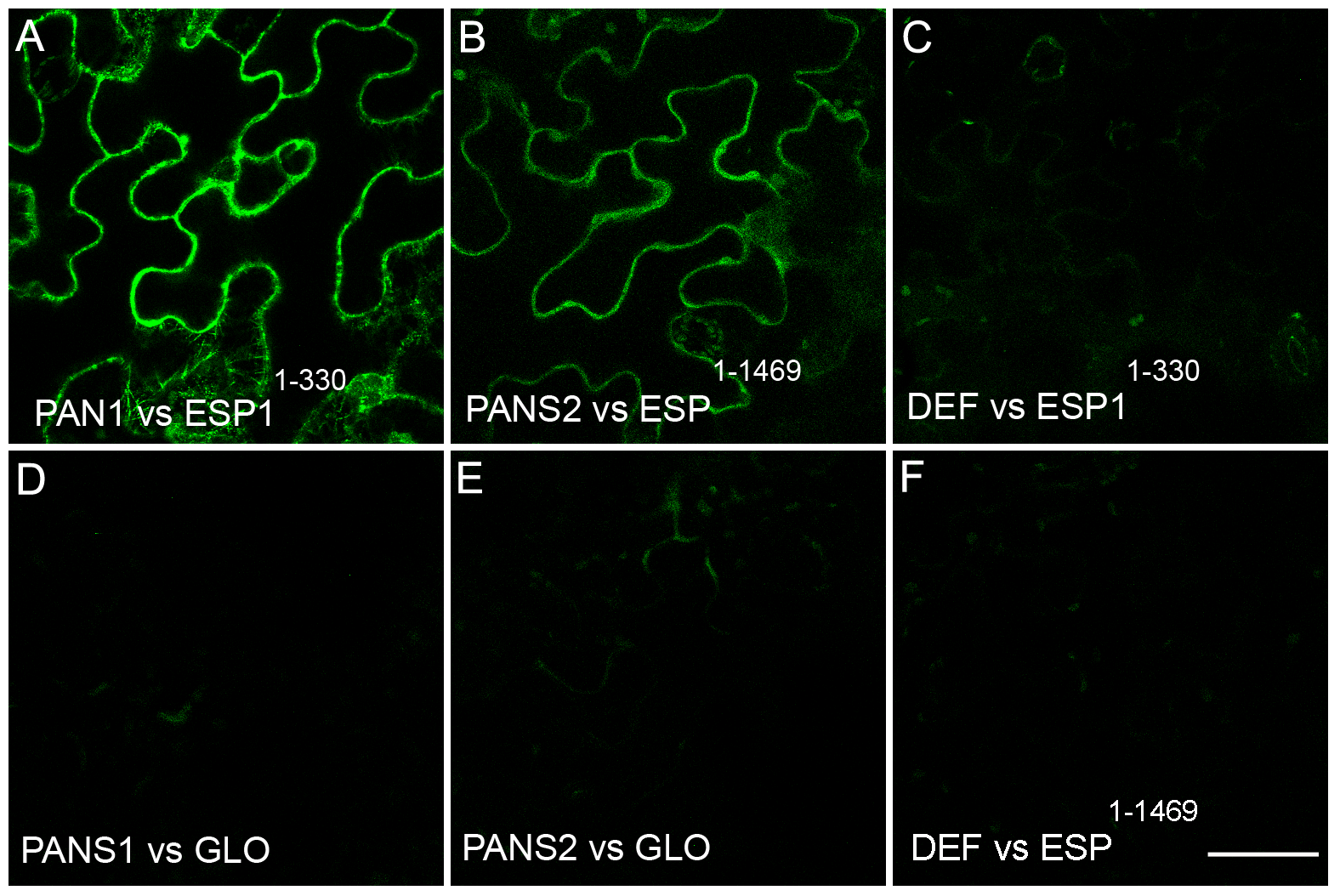

**FIGURE S5. Bimolecular fluorescence complementation (BIFC)**

Protein interactions were tested using BiFC assays in leaf epidermal cells of *Nicotiana benthamiana* plants. (A) Expression of pBIF2-PANS1 and pBIP3-ESP^1-330^  results in a nuclear YFP signal. (B) Expression of pBIF2-PANS2 and pBIP3-ESP^1-1469^  results in a nuclear YFP signal. The four corresponding negative controls (C) pBIF2-DEF/pBIP3-ESP^1-330^, (D) pBIF2-PANS1/pBIP3-GLO, (E) pBIF2-PANS2/pBIP3-GLO and (F) pBIF2-DEF/pBIP3-ESP^1-1469^ do not generate YFP signals. GLOBOSA (GLO) and DEFICIENS (DEF) are two interacting components of the *Anthirrinum majus* MADS box transcription factor. Scale bar=50 µm.
