## Supplementary material for "Patronus is the elusive plant securin, preventing chromosome separation by antagonizing separase": Figure S6

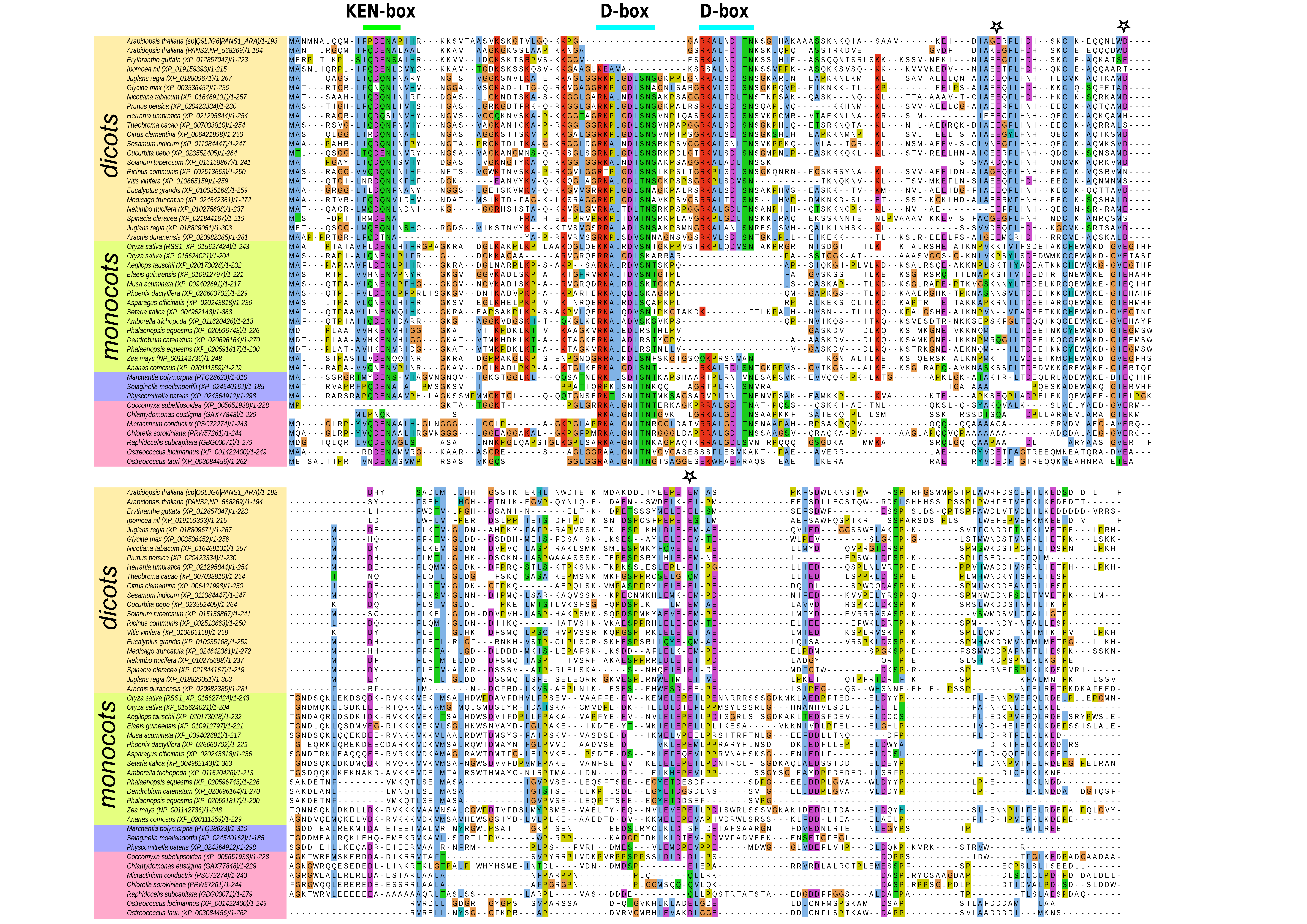

**FIGURE S6. Multiple sequence alignment of PANS1 homologs in plants**

Multiple sequence alignment of the homologues of AtPANS1 in plants built from the full-length sequences retrieved after the PSI-BLAST procedure described in Methods, realigned using MAFFT algorithm and represented using Jalview. Headers are coloured with respect to the clade to which the species belong (i.e. with dicots, monocots, mosses or green algae in yellow, green purple and pink, respectively). For the sake of concision, the columns of the alignment containing gaps in AtPANS1 and OsRSS1 sequences were trimmed.
