## Supplementary material for "Patronus is the elusive plant securin, preventing chromosome separation by antagonizing separase": Table S1

**Table S1. ESP mutations identified in this study**

| Name of original mutant | Number of candidate mutations in the line | Genomic mutation in *ESP* (TAIR10) | Amino acid change in ESP |
| --- | --- | --- | --- |
| *spans25* | 206 | Chr4: 12040670 C>T | S606N |
| *spans41* | 201 | Chr4:12034602 G>A | P1946L |
| *spansD22* | 62 | Chr4:12034107 C>T | A2047T |
| *spansC4* | 389 | Chr4:12033780 G>A | P2156S |
