## Supplementary material for "Patronus is the elusive plant securin, preventing chromosome separation by antagonizing separase": Table S2

| **Table S2.** Homologues detected by PSI-BLAST with the green alga *Ostreococcus lucimarinus* CCE9901 (XP_001422400.1) as the input sequence.  The hits have been ordered by increasing e-values ranging from 4e^-57^ to 1e^-05^ after 10 iterations. | | | | | |
| --- | --- | --- | --- | --- | --- |
| Panicum hallii var. hallii | | (PUZ76441.1) |  | Triticum urartu | (EMS53671.1) |
| Panicum hallii | | (XP_025791674.1) |  | Amborella trichopoda | (XP_011620426.1) |
| Panicum miliaceum | | (RLN07740.1) |  | Brachypodium distachyon | (KQK12869.2) |
| Panicum miliaceum | | (RLM78791.1) |  | Hordeum vulgare subsp. vulgare | (BAK00762.1) |
| Oryza sativa Japonica Group | | (XP_015627424.1) |  | Aegilops tauschii subsp. tauschii | (XP_020182268.1) |
| Oryza brachyantha | | (XP_006647477.1) |  | Oryza sativa Japonica Group | (BAD82353.1) |
| Dichanthelium oligosanthes | | (OEL17437.1) |  | Sorghum bicolor | (XP_021302849.1) |
| Dichanthelium oligosanthes | | (OEL20438.1) |  | Zostera marina | (KMZ64751.1) |
| Oryza brachyantha | | (XP_015689052.1) |  | Dendrobium catenatum | (XP_020696164.1) |
| Zea mays | | (NP_001132520.1) |  | Musa acuminata subsp. malaccensis | (XP_009379917.1) |
| Panicum miliaceum | | (RLM99188.1) |  | Physcomitrella patens | (XP_024364912.1) |
| Zea mays | | (ACG45876.1) |  | Ostreococcus tauri | (XP_003084456.2) |
| Sorghum bicolor | | (XP_002454142.1) |  | Phalaenopsis equestris | (XP_020591817.1) |
| Oryza sativa Indica Group | | (EAY86615.1) |  | Dendrobium catenatum | (PKU68131.1) |
| Phoenix dactylifera | | (XP_008791773.1) |  | Pomacea canaliculata | (XP_025093964.1) |
| Asparagus officinalis | | (XP_020261507.1) |  | Musa acuminata subsp. malaccensis | (XP_018675059.1) |
| Asparagus officinalis | | (XP_020243818.1) |  | Phalaenopsis equestris | (XP_020596743.1) |
| Setaria italica | | (XP_004953025.1) |  | Oryza sativa Indica Group | (EAY76834.1) |
| Phoenix dactylifera | | (XP_026656827.1) |  | Musa acuminata subsp. malaccensis | (XP_018675061.1) |
| Elaeis guineensis | | (XP_010912113.1) |  | Crassostrea virginica | (XP_022340323.1) |
| Zea mays | | (NP_001142736.1) |  | Zostera marina | (KMZ59965.1) |
| Panicum miliaceum | | (RLN29265.1) |  | Physcomitrella patens | (PNR27411.1) |
| Zea mays | | (AQK85829.1) |  | Apostasia shenzhenica | (PKA64779.1) |
| Elaeis guineensis | | (XP_010912797.1) |  | Marchantia polymorpha | (PTQ28623.1) |
| Brachypodium distachyon | | (XP_003575314.1) |  | Micractinium conductrix | (PSC72275.1) |
| Elaeis guineensis | | (XP_019703807.1) |  | Oryza brachyantha | (XP_015690666.1) |
| Musa acuminata subsp. malaccensis | | (XP_009402692.1) |  | Micractinium conductrix | (PSC72274.1) |
| Zea mays | | (PWZ27380.1) |  | Raphidocelis subcapitata | (GBG00071.1) |
| Hordeum vulgare subsp. vulgare | | (BAK07257.1) |  | Chlorella sorokiniana | (PRW57261.1) |
| Panicum hallii var. hallii | | (PUZ66070.1) |  | Chlamydomonas eustigma | (GAX77848.1) |
| Setaria italica | | (XP_004962143.1) |  | Marchantia polymorpha subsp. ruderalis | (OAE27187.1) |
| Ostreococcus lucimarinus CCE9901 | | (XP_001422400.1) |  | Picea sitchensis | (ABK26784.1) |
| Triticum turgidum subsp. durum | | (AIP89948.1) |  | Coccomyxa subellipsoidea C-169 | (XP_005651938.1) |
| Panicum hallii | | (XP_025808982.1) |  | Elaeis guineensis | (XP_010930199.1) |
| Musa acuminata subsp. malaccensis | | (XP_009402691.1) |  | Elaeis guineensis | (XP_010930200.1) |
| Zea mays | | (XP_008647649.1) |  | Acropora digitifera | (XP_015753966.1) |
| Triticum urartu | | (EMS47367.1) |  | Ostreococcus tauri | (OUS44868.1) |
| Aegilops tauschii subsp. tauschii | | (XP_020173028.1) |  | Orbicella faveolata | (XP_020600834.1) |
| Zea mays | | (NP_001158919.1) |  | Crassostrea gigas | (EKC30626.1) |
| Zea mays | | (AQK85830.1) |  | Asparagus officinalis | (ONK57046.1) |
| Apostasia shenzhenica | | (PKA64761.1) |  | Zea mays | (ACG33508.1) |
| Elaeis guineensis | | (XP_019702503.1) |  | Crassostrea gigas | (XP_011441394.1) |
| Phoenix dactylifera | | (XP_026660708.1) |  | Pocillopora damicornis | (RMX43805.1) |
| Musa acuminata subsp. malaccensis | | (XP_009404273.1) |  | Selaginella moellendorffii | (XP_024519951.1) |
| Musa acuminata subsp. malaccensis | | (XP_009404272.1) |  | Selaginella moellendorffii | (XP_024540162.1) |
| Zea mays | | (PWZ17396.1) |  | Zea mays | (ACG37554.1) |
| Asparagus officinalis | | (ONK79989.1) |  | Stylophora pistillata | (XP_022807112.1) |
| Amborella trichopoda | | (ERM98247.1) |  | Orbicella faveolata | (XP_020600847.1) |
| Oryza sativa Japonica Group | | (XP_015624021.1) |  | Lottia gigantea | (XP_009062812.1) |
| Phoenix dactylifera | | (XP_026660705.1) |  | Apostasia shenzhenica | (PKA58368.1) |
| Ananas comosus | | (XP_020111360.1) |  | Saccoglossus kowalevskii | (XP_002742390.1) |
| Phoenix dactylifera | | (XP_026660702.1) |  | Pyrus x bretschneideri | (XP_009363238.1) |
| Ananas comosus | | (OAY72878.1) |  | Lingula anatina | (XP_013412898.1) |
| Asparagus officinalis | | (ONK72469.1) |  | Biomphalaria glabrata | (XP_013082550.1) |
| Ananas comosus | | (XP_020111359.1) |  | Ostreococcus tauri | (OUS44952.1) |
| Mizuhopecten yessoensis | | (XP_021355525.1) |  | Exaiptasia pallida | (XP_020908213.1) |
| Zostera marina | | (KMZ64759.1) |  | Biomphalaria glabrata | (XP_013082549.1) |
| Acropora digitifera | | (XP_015760873.1) |  | Priapulus caudatus | (XP_014665187.1) |
