## Supplementary material for "Patronus is the elusive plant securin, preventing chromosome separation by antagonizing separase": Table S3

**Table S3. Primers used in this study.**

| **Primer name** | **Sequence (5' to 3')** |  |
| --- | --- | --- |
| N570337U | CCGTTAAACACTCTAAGCGC | *pans1-1* genotyping |
| N570337L | ATGCTTCTCCTTGATGCTGG | *pans1-1* genotyping |
| LBSalk2 | GCTTTCTTCCCTTCCTTTCTC | *pans1-1, esp-2, spo11-1-3* genotyping |
| SK2556U | GAGGTCGGCAATAAGGTTAG | *Atsgo1-2* genotyping |
| SK2556L | CCCGATAGACTGTTCACTTG | *Atsgo1-2* genotyping |
| pSKTail1 | TTCTCATCTAAGCCCCCATTTGG | *Atsgo1-2* genotyping |
| SGO2U | TAAGTTATTCCGTTGGCCTAC | *Atsgo2-1* genotyping |
| SGO2L | GATTGCCTTCGTGTACATTGC | *Atsgo2-1* genotyping |
| GABI | ATATTGACCATCATACTCATTGC | *Atsgo2-1* genotyping |
| N537016U | ATGATGTTTTGCATCAGGGAG | *esp-2* genotyping |
| N537016L | TGATAGAAATGGAGGGGAAGG | *esp-2* genortyping |
| N646172U | AATCGGTGAGTCAGGTTTCAG | *spo11-1-3* genotyping |
| N646172L | AATCGGTGAGTCAGGTTTCAG | *spo11-1-3* genotyping |
| esp606U | CTCAAGTATCTAATTGCGACGC | *esp-S606N* genotyping |
| esp606L | GCCTCAGGGCCTGGTAACCG | *esp-S606N* genotyping |
| esp2156U | GAGCACAGTATATACCTAGGCGTG | *esp-P2156S genotyping* |
| esp2156L | GCTTATTGGCTTTTGCCATACTG | *esp-P2156S genotyping* |
| PANS1∆D | CCTGGAGGAGCTCTTAAGGCTGTGAATGATATTAC | *PANS1 mutagenesis* |
| PANS1_XhoI | CTCGAGATGGCGAACATGAACGCTCT | *pDMC1::PANS1 cloning* |
| PANS1_SpeI | ACTAGTGCAGCAAGTTATGAAAAGG | *pDMC1::PANS1 cloning* |
| DMC1_GTW | GGGGACAAGTTTGTACAAAAAAGCAGGCTCAGACCTATATATGTGATGC | *pDMC1::PANS1 cloning* |
| PANS1SpeI_GTW | GGGGACCACTTTGTACAAGAAAGCTGGGTACTAGTGCAGCAAGTTATGAAAAGG | *pDMC1::PANS1 cloning* |
| PANS1cU_GTW | GGGGACAAGTTTGTACAAAAAAGCAGGCTTAATGGCGAACATGAACGCTC | *Y2H & BiFC cloning* |
| PANS1cL_GTW | GGGGACCACTTTGTACAAGAAAGCTGGGTCTCAGAAGAGGTCGTCAGAGTC | *Y2H & BiFC cloning* |
| PANS1cPARTIL_GTW | GGGGACCACTTTGTACAAGAAAGCTGGGTCTCAAGCAGAAGCAATTTGC | *Y2H & BiFC cloning* |
| PANS1cPARTIIU_GTW | GGGGACAAGTTTGTACAAAAAAGCAGGCTTAGCTGTGAAAGAGATTGATATAGC | *Y2H & BiFC cloning* |
| PANS2cU_GTW | GGGGACAAGTTTGTACAAAAAAGCAGGCTTAATGGCCAACACAATTCTTCG | *Y2H & BiFC cloning* |
| PANS2cL_GTW | GGGGACCACTTTGTACAAGAAAGCTGGGTCTCAAGTGGTGTCTTCATCCT | *Y2H & BiFC cloning* |
| CDC20.1cU_GTW | GGGGACAAGTTTGTACAAAAAAGCAGGCTTAATGGATGCAGGTATGAACAAC | *Y2H & BiFC cloning* |
| CDC20.1cL_GTW | GGGGACCACTTTGTACAAGAAAGCTGGGTCTCAACGAATACGATTCACG | *Y2H & BiFC cloning* |
| ESP_1_GTW | GGGGACAAGTTTGTACAAAAAAGCAGGCTTAATGGCGTCCTCTGGCGACGACCTC | *Y2H & BiFC cloning* |
| ESP_2_GTW | GGGGACCACTTTGTACAAGAAAGCTGGGTCGCACAGTACATGTCTCCTGCTGCTCG | *Y2H & BiFC cloning* |
| ESP_4_GTW | GGGGACCACTTTGTACAAGAAAGCTGGGTCGCCTCAGGGCCTGGTAACCG | *Y2H & BiFC cloning* |
| ESP_5_GTW | GGGGACCACTTTGTACAAGAAAGCTGGGTCCTAACGAGATGAAGAAGGAAG | *Y2H & BiFC cloning* |
| **DNA synthesis** |  |  |
| ESP^1466-2180^ | GGGGACAAGTTTGTACAAAAAAGCAGGCTCCACCATGGCGTCCTCTGGCGACGACCTCCGTCTTCTCTCTCTCATCGACGTCGGCGATAATGTCTTCTCTTCCTTCTCCGATTACCTTAAACCTTTCTCTACTCTATCCACTTCCCGGAAAAAACAAGACCGAGCAACCACGATTCGAGCCCTAGCTAAGCAATTCCTTCCTTTCCTCAACAAATCGATCTCTCTCCTACCGAAACGTCTCTCCGTCGCGAATTCTGATAAGGAAGCTCGTGAATCAGCATTAGATCTGTTCCGTGCTTATGAACTTTGTTTGGACTGTTTAGAATTGGTCTCTGCTCAATTAGCTTGTAAGCCTCACACTGTTCAATCTCAGAGGCTAAGGATGATTCATTGTCTCGATGTTTGGGGTTTGTATGAGAATGTGTATACTGAAGCGTTTAAGGTTCTGGAGAAGCTAAGAGGCTCTGATAGCAAATCCCGGAAGTCCCGGTTGTTGCCGGAGGTTCAGGATGGAGATGCGGAGATGGCTTTAGTTGTTGTTGATGCTGTGGCGGCCATCTTTAGGGCTGTGGCAATGAGTCAGCAGTTAGATGATAAGCGGTATCGGAAAGTTCTTCTTTTGCTTGAGGAAGTCGGAGGCTGGTTAAGGGTCTTAGATGCTAAGGTGTATGAAAAGTTGCACAGAGCGATGGTGACAAGTATGGGTAAATGTGCTGTTTCTCTCGTTAGAGAAGCTGAGCGTTTCAATGGGGATTTAGTGATTTCTTTCTGCGACTTAACTGTGAAGGAACATTATAAATCTGCATTATCAAAAGATCGAGTTTATAAGTTTGCTCGCGAGGTGCTTTCTGTTCTGTTTGGATTTAAGGATAGAAAAATGTCAGTGACCATTGACATTTCAATGTCTGTGTTGCGCAGCTTATCTTGCCAATTTGAGGATGAAAGTAATGAGAATTTAATGGAGTTTTTTGACCTAGTCGATTATTGCGCTCACAAGTTTCGAGCAGCAGGAGACATGTACTGTGCAAAAGTTTCAAAAAAATTGAATGAGATGGCAGCTATTTTCGTAGAGGCAATACCCCAACTTAATTTGGTTCTTAGACTGTATTCGACCGGGCTGTCCATCACGGTTTGCAATTCCAAGCTCGGAGAAATTAAGCTAGAAGATTCAACAGATGATTGGAAGATTCAAGCTATGTTTGATGATGATGCTAGATGGCAAAGCTTAGTTTCTTTACTCGGCATGGTTGATAGTTACTCAGGTGATGAGGGCAATCAGACTGGTTCATCATCGATTGGTGGGCATAGGAACTACAACAATAAAACACATGATAGCTGTAAAGACAGGAACAAGATTACTTGTTGGCCACAATACGTAGATGCCTTGAAATTCTTGTGCCAACCGCTTGCAGACTTTATATATTCTGTGAAAAGAAAGATTGTGCTGGAGACAGAAATGTCATGCGCCTCTGCTCATCTAATTACTATTCACGATGCCTTTCTCCAGTTTTGTGATGGCTGTCTCTTCCTTCAGAGGTGCACGTCTGACAAGGGAGATAGAGAAATTGCCAATAACAAAGCTTTCTTAAACGCGGCTATGGGTGCTTTCATTGTCTCATTGAGAACCCAACTTAAATTGGAGATAAGTGCCCATCTAGTTGAGGATGTAATTGGAAGCCCATGGATCCAATCTCAAGAACTCAAGTATCTAATTGCGACGCTGTACAATATTGGTATCGTTTTGTACAGAAATAAGGAGCTAAACAAGGCTTGTGAGGCGCTTAAGCTGTGCTCCAAGGTATCATGGAGATGTGTTGAACTACATTGTCATATGTTTGTAAATCAGTCTAGTTCATCTGATAATGACTTGTCAGAAGATGCCATTATGGATTTTGTGGGCGAAGCATGTAATAGGTGTGCATTCTACCTGGATATACTCCAGAAATGTAGCAGACGTAAGATTAGACAGAATATTGTTCATATTCTAGAAAATTGGTTGTCAGCTGAACATCTGATTAGACGGTTACCAGGCCCTGAGGCAATTGTAAAGCAATGGGTTAAGATAGAGCGGGAATGTCACACGGATCTGGATGCGGCAGGTTCTTGTACAACTTTGTACTCATTGTTATCATCTTCTCAAAAAAAGTCAAAGCGGGGTATCGGAAAAATCCTTGAGCAGGAGCTTCTTGCCTATGACAGAGTGTTACCTTTAAGGTCAAATCTAGGTCAACAAACGCGAATCAAGATAGCAGATATCCTCTTGAAGGATGTTTATGTCACTGAGGATATGCATATAGAGAGAGCAAGAATTTTGATATGGAAAGCACGAATGACAAGGACATCTGGAACTGAACATATAACCGAGTGCATTTGTTTTCTGTCTGAAGCAATCTCTATATTGGGTGAGTTACATCATGGGCCAAATGAAGAGGGTTCTCCATCTTCTCACATGTTACCTATTGCGTATTGCTTGAGAGCTTTTTGTACCCAGGAAGCTGACCCAAACTCGAAGAAAGTCTTTCAAGATATTAGCACTTCACTAAATCTCTGGCTAAGGATTCTTAGCCTAGATGATAGTGGAGACAGTCTACCAACAGAGAATATAATTCCGTTACTGTATAATATGATTGATTTGATGTCAGTGAAGGGATGTACAGAGCTCCATCATCATATATACCAACTGATTTTCAGATTGTTTAAATGGAAGAATGTCAAATTGGAGGTCTGCCTTGCAATGCTGTGGGAATGTAGAAGGCTTAGTCATGCCTTATGCCCTTCCCCCATAAGTGATGCCTTTATCCAGACCTTATCAGAGAATTGTGCTGACAAATCCACATGCATTGATTTCTGGATGGATTGCCTGAAAGATTCAAAAGCAAAGTTAATTGGGTTCCAGCAGAACTTCCATGATTTACATAATAAGGATGAAGGTCCCTTTCAATCAGACATCACAATTGACGATATTAAAGATGCAGCATCAGAACTCATCTCCAGTGCATCACTCTCTGGTAATTCATCTTTTCAGCTGCATATCTCTATTATGATCTCTGTGAAAGGCTCATATCATTTGGAAAACTTTCTGAGGCTCTTTCATATGCAAAAGAAGCCTATAGGATAAGGACTCTTATATTTCAAGATAAATTTAAGTATACAGCTGAGAAGCATATCGAAAAACACAATGAAGATGGGAAAATATCAGAAATACGGACTTTTAGCATAAAAAATTTCCAAGTCTACAGATTGTTAGCAACTGATTTTTGGCCATGTGGAAATTTTCTCTGGGACATCAATCGCTGCTACCTAAGTCCTTGGAGTGTACTCCAATGTTATTTGGAAAGCACCCTTCAGGTTGGAATTCTGAATGAGCTCATAGGGAATGGACTGGAGGCAGAAACCATTCTATCATGGGGGAAAGCCTTTTCATGCTCGCAAAGTCTATTCCCTTTTGTAGTTGCATTTTCTTCTGCCTTAGGAAATCTATACCACAAAAAGCAGTGTCTGGATCTGGCAGAAAAGGAACTCCAAAATGCAAAAGAGATATTAATTGCTAATCAGCGAGATTTTTCTTGCGTAAAGTGCAAACTGAAGTTGGAAGTTACATTGGATAAACAACTTGGAGACATATCTCGGAAACAAATTGATAGAGTTTCACAGACAGATGGATTCTTGCATGCTGAAAGCTTGTTTAGTGCTGCCCTGGGAAAATTCTGTTGCTCAGCATGGAAAAGCTGCATTAGATCACATGGGGAAGAAATTGCTGAGGAGATAGTGATTGATAGAAATGGAGGGGAAGGTTTAGGACATAATTCAAGTAAAACAAAGCTCAGTATAAAGGAGCCACCAGGAAACAGAGGCTCCAGGAGGGGTGGAAGAGCTAACAAAACTTGTTTGTCAAAGGATCAGGATTTGATATCTGAGCCAACATCAAGGTTGACTCGATCTATGCGTCACTCACTTAGAGAGCAATGCCAAAACCGTTCTAATGTGCCTGAAGTTGTTTCAAAGAAGCCTAATTTATGTGATAGATCTGTTGGTTCTAGGGGTGAAAGGGTTTTGTTGGACACGAGTAATGCACTCCCTGGCTTTTGTATTTGCTACAAAGAAAAACGTCAGCAGTGCCTCTCGGAAGAAGTAACAGAATCTGGGTCTCTCAACAATTTAGTAAGTTTGAAATGGGAACTCTGCCATAGGAAGCTTGCGTCCTCAATACTTGTTAGCCTTGGAAAATGTTTGGGAGACTCTGGTAGAATTCATCTAGCTCATGAGGCTTTACTGCATAGTATCTCTGTTTTATTCAAAAGCACCTGGTCCAGCCACAACCAGCCTTCTGTCAGTCAGTTGCTGGAATTTATCGGGAAAGAAGTTACAAGAGATGTATTTGCCGTTGATCGTGCAATAATATTATACAACTTGTGCTGGTTGAATTTGCGGAATTACCACTGAGACCCAGCTTTCTTGTACAAAGTGGTCCCC | |
