## Supplementary material for "Patronus is the elusive plant securin, preventing chromosome separation by antagonizing separase": Figure S3

```
Arabidopsis_thaliana       1    MASSGDDL-RLLSLIDVGDNVFSSFSDYLKPFSTLSTSRKK--------QDRAT----------------------------------------------------------------------------
Oryza_sativa               1    LA--------------------------------------------------------------------------------------------------------------------------------
Chlamydomonas              1    MPAAGDSQ-A-----------------------ALATVEES--SDVRKLQDALGTL-----------------------------------------------------------HELLAPLLALRDGSV
Homo_Sapiens               1    LGSLL-EL-A-----------------------ELA-----------CDGYLVS----------------------------------------------------------------------------
Caenorhabditis_elegans     1    MKIT------------------------------------------------------------------------------------------------------------------------------
Saccharomyces_cerevisiae   1    MMVKQEEPLN-----------------------EISP-NTP-----------------------------------------------------------------------------------------
Schizosaccharomyces_pombe  1    NLAKQWA--------------------------ELSTTDKEKTRRMFCTPSRLNTAHRPEVFYLLECCTYILEQMQVVTKNTSHLYDCIRSGVSICNRLLDMEIFEPAISLLMKTHKNLIILLTYRDH--

Arabidopsis_thaliana       45   -TIRALAKQFLPFLN-KSISLLPKRLSVANSDKEARESALDLFRAYELCLDCLELVSAQLACKPHTVQSQRLRMIHCLDVWGLYENVYTEAFKVLEKLRG-------SDS--------------------
Oryza_sativa               2    -------KRFLPFLC-RALQVLPLLLRPNP-SSGDAGCTDELLDVYALLLDCLAVISACLAGKPYSVLLQRGRFVCCLESRGHYARAEADAAATLDSLRS-------VLSV-------PTAS--------
Chlamydomonas              46   DALRPIAKTCGPVVT-RLVKSTGCRL--G-SELRGSQVADILGDAACLGLNALDCLRPVIKASGHELDTQRYGFIRKLAALGLHERALEQ-GRVLHAAV-------------------------------
Homo_Sapiens               18   ------TPQRPPLYL-ER--ILFVLLRNA-AAQGSPEATLRLA---QPLHACLVQCSREAAPQDYEAV-ARGSFSLLWKGAEALLERRAAFAARLKALSFLVLLEDESTPCEVPHFASPTACRAVAAHQL
Caenorhabditis_elegans     4    --------------------------------------------------------NKSVDKQHI---------------------------EKLDELRK-------NVSCTVIGFAEQTAELQQEISEL
Saccharomyces_cerevisiae   17   --------MTSK------------------------------S---YLLNDTLSKVHHSGQTRPLTSVL-------------------------------------------------------------
Schizosaccharomyces_pombe  103  DAI-PTATLLNPTLDVSEIQLESCLF--V-PMVPASYFLNIGTIVVTFQLNVLRCLS-LSQINGLSLNTINN-------------LQ-SE---------------------------------DGPFQWI

Arabidopsis_thaliana       146  ----------------KSRKS------RLLPEV-QDG---------DAEMALV-----VV--DAVAA--IFRAVAM--------S-QQLDDK-------RYRKVLLLLE---------EVGG-WLRVLDA
Oryza_sativa               101  ----------------KSRRAATASFASLLPDPGISGEAGA-----DPEVTIL-----AI--ELTVC--FANCASK--------C-KVKEAA-------PYERVVSLVD---------QLQP-WLRILAE
Chlamydomonas              139  -AAHP-------------------------------SNAGA-----PAELQVA-----AVA-NLLLC--YGELAQR--------D-PYRSEELWQGIGAPVESLLDLAREADKEDCARRLEP-VLR----
Homo_Sapiens               135  FDASGHGLNEADADFLDDLLS------RHVIRA-LVGERGSSSGLLSPQRALC-----LLELTLEHC--RRFCWSR-----------------------HHDKAISAVE---------KAHS-YLR----
Caenorhabditis_elegans     45   FIAEFGVNGPIDMNSLSKL---------------------------------------------------------------------------------------------------------AR----
Saccharomyces_cerevisiae   45   -----------------------------------SGDASS-----N-SIGIL-----AM--HNNIIRDFTKIASNNIDLAIEDITTVD--H-------SLNSIYSLLK---------SHHM-WGHINS-
Schizosaccharomyces_pombe  181  ERSF------------------------------------------PSQVQLANSRREILA-RLLTR--FS-MIQN---------NALQSFKLLILSIALWLNILSSQRADDKEFDVNQLETRILQ----

Arabidopsis_thaliana       210  KVYEKLHRAMVTSMGKCAVSLVREAERFNGDLVISF--CDL----------------------TVKEHYKSALSKDRV--YKFAREVLSV-------------LFGFKDRKMSV--------------TI
Oryza_sativa               176  DVSRKYLTLLVNALSRCAILLVAEYSVFNTNLVCEF--CRA----------------------TLGECMK-AQTIERL--PAVARKICSS-------------VDVSWGGGTQL--------------LL
Chlamydomonas              210  ----------------------------CLGKAVGALF-SA----------------------SLSTHLEA-STVA-------------------------------------AQLAALVACGDVCGQEQ
Homo_Sapiens               213  ----------------------------NTNLAPSLQLCQL----------------------GVKLL--Q-V--------------------GEEGPQAVAKLLIKASAVLSK--------------SM
Caenorhabditis_elegans     65   ----------------------------ITSYYASSEYFQGLAKYQRTACKMFITWQTLRKEAME-----C-RSKDREIFASIPAKLCFFYFYNGELCRAVVC---------------------------
Saccharomyces_cerevisiae   107  -TVKQHLMIIVKLINNNALGLASSEIIFLFNETNLF--QAH----------------------SLKNI--------------------------------------------------------------
Schizosaccharomyces_pombe  251  ------------LFSK-VVQL-CKSEDIEGSI--LN--KDM----------------------TQLHHLLENLSKESR--LHILLQLSQL-------------YYKYNDFQLSA--------------A-

Arabidopsis_thaliana       287  DISMSVLRSL--SCQFEDESNEN----------LMEFFD-LVDYCAHKFRAAGDMYC--AKVSKKLNEMAAIFVE-------AIPQLNLVLRL---YSTGLSITVCNSKLGEIKLEDSTDDWKIQAMFDD
Oryza_sativa               252  DVLKTVVGSA--ACLKADLP-RA----------VNGLVE-FVAYFSRSFVSSNWDLS--VGAAELIYEQGGYFSEV-----SSTPATASVLIL---YAIGLYCSA-------QQIENRERPHKSTDFLND
Chlamydomonas              252  AVK-----------------------------------------QLLVDDAAE-------------------AAEE-----ARVALLEALLEQ---LEM-----------------HGALPDVLCCSLHK
Homo_Sapiens               257  EAPSPPLRALYESCQFFLSGLERGTKRRYRLDAILSLFAFLGGYCSLLQQLRDDGVY--GGSSKQQQSFLQMYFQG-------LHLYTVVVYD---FAQ-----------------GCQ-----------
Caenorhabditis_elegans     134  ---------------------------------LLD-YIDLSDDTLAKEAALRWL-------------------MF-------LGETELIEKKLKTWKM-----------------DKS-----------
Saccharomyces_cerevisiae   150  --------------------------------------------------------------------LLADFSTW-----NDYYLSNLKILA---LQIILKR----------KLVDEYLPHILELFSHD
Schizosaccharomyces_pombe  311  ----YVIRGY--SLSFEDISFKL----------KFLLFSF-------RLSIHDNSICFPFNLIQELSSLQQLFVENALPYSEALHLLDSIERS---FRL-----------------FN-----DSTVFD-

Arabidopsis_thaliana       392  DAR--------WQSLVSLLGMVD---------SYSGD-EGNQTGSSSIGGHRNYNNKTHD-----SCKDRNKITCWPQYVDALKFLCQPLADFIYSVKRKIVLET---EMSCA----S------AHLITI
Oryza_sativa               351  EKH--------LQTLKSALATLAHLFCFANGRSIPLDTMGKASSSSMQPGHSNKKNSLSH-----S----DDHISFVAYLDSLEFLCKILSQYVNAVWKNFS-EG---ITPNY----S------INMTYV
Chlamydomonas              297  LALGLVQGRISA---------------------------------LTVAAHR-----PHPLAAIVQAALSVPACAPVMAPAERQKQQHPLAAAL--------ARMPQAASAMS----E------EQKLSL
Homo_Sapiens               346  ----------------------------------------------------------------------IVD---------------------------------------------------------
Caenorhabditis_elegans     176  ----------------------------------------------------------------------SKD---------------------------------------------------------
Saccharomyces_cerevisiae   195  KRY--------LL---------------------------------KDPNLK-----AHALTKIVLSFFSVTTSCK--VLFGLKFL-QYIKQFKLPF-KKFISNI---TVECF----S------KNLLHK
Schizosaccharomyces_pombe  392  -----------------------------------------------------------------------------DTVFALN-ISEILSWILSSVVRDILVED---ELLNLQLKIRKFLMFTFHIIRS

Arabidopsis_thaliana       485  ----------------------HDA---FLQFCDGCLFLQ-----------------------------------RCTS-DKGDRE--IANNKAFLNAAMGAFIVSLRT----------------QLKLE
Oryza_sativa               449  ----------------------LTA---LHQFINSSIAAY-----------------------------------SCTEMPEGDKDKQHEQHGTLLRALVSAMKVSFIT----------------NEGIQ
Chlamydomonas              370  ----------------------SDR---WLRWC--CLDVL-----------------------------------RVSVA-GVLTH--LLNNPQRAGGHLPAVAVCMRALAADSLQLLHQSP--GRASTV
Homo_Sapiens               349  --------------------------------------------------------------------------------------------------------------LADLTQLVDSCKSTVVWMLE
Caenorhabditis_elegans     179  --------------------------------------------------------------------------------------------------------------MFSATEFAMNY--------L
Saccharomyces_cerevisiae   262  NYLEMGPNKIYLNSFYLSYSMLYDG---LDKIMLLDILSYEETTEVQRAIKSKKEFNEYCNMSENRLLWSCISVDDLNV---ILEN--ATNFLQNKGKHISATLK-------------------------
Schizosaccharomyces_pombe  441  ----------------------FSELTKFQSSLEGCLNLA-----------------------------------AYYE-D--------AEFPQKLSNHLYNLC-----VKSSNVN-YARECISLSIKIA

Arabidopsis_thaliana       537  I---------------------------------SAHLVEDVI---------------------------GSPWIQSQELKYLIATLYNIGIVLYRNKELNKACEALKLCSKVSWRCVELHCHMF-----
Oryza_sativa               504  K---------------------------------SVSFIKCAI---------------------------SSTWIKLDEIKFLMSSLGNIGVTLYNIGHLDEAPKALELCCQTVWVYARLSYHRL-----
Chlamydomonas              434  VPSGCSAATLACRLMAAHRTSTGQLAGLHEDVAAGAKLLRDLLSASSQRPQQQPQSTSMNDEEQDGPCSGAADAVTPGMLQSLSASMGNAGLDLLQAQHVAPAAELLQLSADLALRRAQALH-G------
Homo_Sapiens               370  ALEGL------------------------------------------------------------------SGQELTDHMGMTASYTSNLAYSFYSHKLYAEACAISEPLCQHLGLVKPGTYPEVPPE--
Caenorhabditis_elegans     192  KKSEY------------------------------------------------------------------RVEMLEKLMKLRDKVKSD----------PTRSFSRYELASYVSWLCSTLSN--------
Saccharomyces_cerevisiae   358  -------------------------------------------------------------------------------------------------------------CLVCLWSTIRL-EG-LPKNKD
Schizosaccharomyces_pombe  500  V---------------------------------SHKLT--------------------------------------NDETYLLKILKNFQLRYHDSLQLQEKCDVLHTT----FNQLDLYVGTT-----

				*  esp1-S606N
Arabidopsis_thaliana       601  -----------VNQ-SS-SSDNDLSEDAIMD-FVGEACNRCAFYLDILQKCSRRKIRQNIVHILENWL---SA-----EH-------LIR-RLP--GPE------AI-VKQWVKIERE------------
Oryza_sativa               568  -----------SASQDEQRIIEDIPKDTLKD-ISMDAFAKITKMVDILHRCGVKIIPDIIVKSLSELL---AN-----DS-------TSE-FLN--SSL------VL-IKLWVKITHK------------
Chlamydomonas              556  ---------A-AAGT---MADPATTAAAAAD--VAAVVKKCKAH----------VSALQQVSDLEGVM---AA-----SAAYLAQLHGLC-EAH--VAL------PL-VKLYVRASLTRAHNAAAAAHTA
Homo_Sapiens               432  KLHRC--FRL-QVES---LKKLGKQ---------AQGCKMVILW-----------------------LAALQP-----CS--------------------PEHMAEP-VTFWVRVKMD------------
Caenorhabditis_elegans     237  --------------------VPV-----------GSALRECEFP-------------DRVSHIQEAAL---KS-----DS-------LVRNRIP--GLASSQFDNSVNASIWPFL---------------
Saccharomyces_cerevisiae   378  ILRQFDCTVIYINS-NI-KSINDESAAALLSELLGVLSEICIDYK----------EPKRLSNIISVLF---NA-----SV-------LFKSH-------------S------------------------
Schizosaccharomyces_pombe  549  ---------------------------------------SV----------GKSSVLDNILKRIFN------SLTSINDS-------NIE-KLLESISY------SL-LKLFFKCANE------------

Arabidopsis_thaliana       680  -----------------------------CHTDLDA------------------------------AGSCTTLYSLL----SSSQKKSKRGIGKILEQELLAYD------------------------RV
Oryza_sativa               649  -----------------------------DAKDDES------------------------------VDSAPLLYHSL---MGCTPPLPTKLVGLILEQELLAYA------------------------LV
Chlamydomonas              644  ACSSAAEPAHSAPSTPTAPAAPGPAAPCKARRAGRARAQEVQPKAAAKAPRGRTRLALDADTNTDAGDGAPVKSISLQSAAGA-PPVPMPAESLSVLQAMVSAVESGAAPAGLLGCLDVYAAPELELAAA
Homo_Sapiens               485  -----------------------------AARAGDK------------------------------ELQLKTLRDSL---SGW-DP---ETLALLLREELQAYK------------------------AV
Caenorhabditis_elegans     291  --------------------------------------------------------------------------DGH---QED-----------------------------------------------
Saccharomyces_cerevisiae   443  ----------------------------------------------------------------------------------------------------------------------------------
Schizosaccharomyces_pombe  597  -----------------------------GSRYNAS------------------------------A--------------A----LSFKLSLMLHEK--------------------------------

Arabidopsis_thaliana       724  LPLRSNL--GQQTRIKIADILLKDV-YVTEDMHIERARILIWKAR-----MTRTSGTEHITE--CICFLSEAISILGELHHGPNEEGSPSSHMLPIAYCLR-AFCTQEADPNS-----------------
Oryza_sativa               694  ESRGTMF--CVEMQKRITNILLNKI-YCSKEYYLERSRVLVRKAR-----VLRTCGVQSISS--CLESLSEAISLLRDIPLDSSQGNAPAIHQLAIAYCLH-AHCAQEANLGA-----------------
Chlamydomonas              773  E---CRRMADPRLQQAHADAVVGALFRCAERRGAEMAPVLRARALLLRALLGCPNGAAAVAQ--AGRDLQEACELLD-----------------------------------------------------
Homo_Sapiens               526  R---ADT--GQERFNIICDLLELSP-EETPAGAWARATHLVELAQ-----VLCYHDFTQQTNCSALDAIREALQLLDSVRPEA-QARDQLLDDKAQALLWL-YICTLEAKMQEGIERDRRAQAPGNLEEF
Caenorhabditis_elegans     297  -----------------------------SNYYVHIGSTIAWHFE-----MRRECALVNVTTAQTRDSMSAMILNLR-----------------------------------------------------
Saccharomyces_cerevisiae   443  ------------FLLKTANLEISNV-LISNDSKTSHRTILKFEKF-----ISSAQS---------------------------------AQKKIEIFSCLFNVYCMLRND-TL-----------------
Schizosaccharomyces_pombe  618  ----------------------------------------------------------------------------------------------------------------------------------

Arabidopsis_thaliana       825  -----------------------KKV-------------------FQDI-----STSLNLWLRILSLDD----SGDSLPTENII------------------PLLYNMIDLMSVKGCTELHH--------
Oryza_sativa               795  -----------------------EVI-------------------FDSA-----QNVFGLWSKIKTFGYYSPGMISQQPSENLV------------------PLLCSLVDLLAMKGCFELQF--------
Chlamydomonas              844  ----------------------------------------------------------------------------------------------------------------------------------
Homo_Sapiens               643  EVNDLNYEDKLQEDRFLYSNIAFNLA-------------------ADAAQSKCLDQALALWKELLTKGQAP-AVRCLQQTAASL------------------QILAALYQLVAK-PMQAL----------
Caenorhabditis_elegans     340  ------------------------VA-------------------LKS------ASFFRVLQTTNTLAYYS-SIIEEAGSEKNA------------------K---------------------------
Saccharomyces_cerevisiae   504  -----------------------SFVFDFCQNAFIHCFTRLKITKFIEF-----SNSSEIMLSVLYGN----SSIENIPSENWS------------------QLS-RMI-FCSLRGIFDLDPLELNNTFD
Schizosaccharomyces_pombe  618  --------------------------------------------------------------------------------EEVLLLKTNVSCVLANHGYNDIKFE-EMV-LCVIKGD--QNLLEHNSNNN

Arabidopsis_thaliana       878  -----HIYQLIFRL----FKWKNVKLEVC---------LA---------------------MLWECRRLSHALCPSPISDAFIQTLSENCADKSTCIDFWMDCLKDSKAKLIGF-QQNFHDLHN------
Oryza_sativa               852  -----DLCKLMIII----WKQENLPPEKL---------FS---------------------MLFTNGRLHHACCHLPMDQQFISIAEHHLDVDCHSTEFWRNCFKGDHPSLCMF-LQRLWPIDS------
Chlamydomonas              844  ----------------------------------------------------------------------------------------------------------------------------------
Homo_Sapiens               723  -----EVLLLLRIV----SE---RLKDHSKAAGSSCHITQ---------------------LLL-----------------------------------TLGCPSYAQ----LHLEEAASSLKHLDQTTD
Caenorhabditis_elegans     375  ----------------------------------------------------------------------------------------------------------------------------------
Saccharomyces_cerevisiae   583  KLHLLNKYELLIRI----VYLLNLDMSKH-----------------------------------------------------------------------------------------------------
Schizosaccharomyces_pombe  665  AKLALNE-SLLCSWENLLCYRRAEDDSRI---------LTIIESWTIFISRFSSVISRCSFT--------------------------------------------------------------------

Arabidopsis_thaliana       962  ----------------------------------------------------------KDEGPFQS----DITIDDIKDAA-SELISSASLSGNSS----------------------------------
Oryza_sativa               936  ---FIST---------------------------------------------------TCEPSFRREFGFGGSVHEVDSVA-SSLVSDATVNDQST----------------------------------
Chlamydomonas              844  -------------GQL---------------------------------QEQGLGHIAARPGAGGG----DDSEDDAKEQAGGQALSAAALGLLDSAALAHAQLGLWLAQQRMATSQ-------------
Homo_Sapiens               782  TYLLLSLTCDLLRSQLYWTHQKVTKGVSLLLSVLRDPALQKSSKAWYLLRVQVLQLVAAYLSLPSN----NLS-HSLWEQLCAQGWQTPEIALIDSHKLLRSIILLLMGSDILSTQKAAVETSFLDYGEN
Caenorhabditis_elegans     375  ----------------------------------------------------------------------------------------------------------------------------------
Saccharomyces_cerevisiae   607  ----------------------------------------------------------------------------------------------------------------LTTNLSKITKLYIN---K
Schizosaccharomyces_pombe  716  ----------------------------------------------------------------------------------------------------------------------------------

Arabidopsis_thaliana       995  ----------------------------------------------------------------------------------------------------------------------------------
Oryza_sativa               977  ----------------------------------------------------------------------------------------------------------------------------------
Chlamydomonas              911  ------------AC------DE-AMEHV---GKAARLWARILACRTCHSSVSQLRMPQASLQALLQAHHLVHLLAGSFPREASQLR-----------RVTP-RLV-----SWLAQQVKLREQH-------
Homo_Sapiens               907  LVQKWQVLSEVLSC------SEKLVCHL---GRLGSVSEAKAFCL--------------------EALKLTTKLQIP-RQCALFLVLKGELELAR-NDIDLCQSD-----LQQVL-FLLESCTEFGGVTQ
Caenorhabditis_elegans     375  ----------------------------------------------------------------------------------------------------------------------------------
Saccharomyces_cerevisiae   623  WLQKSDEKAERISSFEMDF-VKMLLCYLNF-NNFDKLSIELSLCI--------------------KSKEKYYSSIVP-YA-DN-YLLEAYLSLYMIDDALMMKNQ-----LQKTM-NLST--AKIEQALL
Schizosaccharomyces_pombe  716  -------------DFEINSILNFFFCFLHTVEPSGKLTFELAFL--------------------------EIFYE----L-FN-CLLHLQFSKYLVIIGTLLSDKYMTLGFSGKA-HLFY--TKCYSYLR

Arabidopsis_thaliana       995  ----------------------------------------------------------------------------------------------------------------------------------
Oryza_sativa               977  ----------------------------------------------------------------------------------------------------------------------------------
Chlamydomonas              995  ------------------VYLLLTALPDSTCATWAMAAAPAASLAEAAAE--AVEAPSTKLGELLELMQPMR-----------AAPLLDHQALGSAAAGCAA-ELERGGFLR-HGPAPLLLRCRCHQSAA
Homo_Sapiens               1000 HLDSVKKVHLQKGKQQA-QVPCPPQLPEEELFLRGPAL----ELVATVAKEPGPIAPSTNSSPVLK-TKPQP-----------IPNFLSHSPTCDCSLCA-S-PVLTAVCLRWVLVTAGVRLAMGHQAQG
Caenorhabditis_elegans     375  ---------------LM-RVSCVNLLSSNPIIVRCSTP----KETGATSRAHTPMAGSSVSEKQNT-MRPDLADLLGDLELLDEQSFHPITRSCVCNVCT-IYPLHSSFAAEY----------MMSYAIH
Saccharomyces_cerevisiae   720  HASSLINVHLWDSDLTAFQIYFGKTLPA---------------------------------------MKPEL-----------FDINNDHN-------------LPMSLYIK-----VILLNIKIFNESA
Schizosaccharomyces_pombe  799  QCKSSPFINFWNVSYGKYLILTGNTDKGI--------------------------------------LQLKK-----------YSLSSEE-------------DFNSNGLSR-----TVSL---------

Arabidopsis_thaliana       995  ----FAAAYLYYDLCERLISFGKLSEALSYAKEAYRIRTLIFQDK--------------------------------------------------------FKYTAEKHIEKHNEDGKISEIRTFSIKNF
Oryza_sativa               977  ----FLAGYLYFDLSERLLSRGELFQAFSYGKEALHLRKKLLRKKFK--FN-------------FG-KFTSGEAQCSGG--QNSVSLEAWGSTITEIWPDST----------------------------
Chlamydomonas              1093 AAYLR---------------AGDTANAYVQAQEALRLTCGLFALVDT--PDPSPDSAVPSHA-ASA-ASASGRGAGAGA--AKPTPAEAAAAAVQSGAATSA----------------------------
Homo_Sapiens               1111 LDLLQVVLKGCPEAAERLTQALQASL--NHK---------------------TPPSLVPSL-LDEILAQAYTLLALEGL--NQPSN-ESLQKVLQSGLKFV-----------------------------
Caenorhabditis_elegans     474  SDFSQLSIKHFNDEFARIRERGMSSQVLMHRD--------------------SSVRPRPNIIQNEIFGMCVIRWLTKKLDSKESAD-EDTMEIFNNALKIV-----------------------------
Saccharomyces_cerevisiae   782  K---------------LNIKAGNVISAVIDCRKAQNLALSLLKKKNK-----------------------------------------------------------------------------------
Schizosaccharomyces_pombe  852  ----NLLLYERIQLSDALFQLGYTTVSLGFIMQNLKVIKGLFSKSSKEHFN-------------GG----------------------------------------------------------------

Arabidopsis_thaliana       1066 QVYRLLATDFWPCGNFLWDIN---------------------------------------------RCYLSPWSVLQCYLESTLQVGILNELIGNGLEAETILSWGKAFSCSQSLFP--FVVA-------
Oryza_sativa               1057 ---------------RSTGTR---------------------------------------------DSFLTPWTVLQCYLDSILQVALLHELIGNGAEAEVLLRTGKDISQFQGLPV--FGVL-------
Chlamydomonas              1173 ---------------HDYAVRDEDNEPKDEFEDLGRRPGQQQQAAASGDPQREVVGVEGSKQSALGVSAGLGWAVAAAHLASLYQAARVFEASGCVEDAMCLWRECARAAAAFGVRS--LQAL-------
Homo_Sapiens               1184 -------------------------------------------------------------------------------------AARIPH--------LEPWRAS--LLLIWALTK--LGGL-------
Caenorhabditis_elegans     553  -------------------------------------------------------------------------------------R----Y--------LQQR--T--TDMILAVTQ--LGRQ-------
Saccharomyces_cerevisiae   813  ------------------------------------------------------------------LSQGSRLALLKSLSFSFFQLIKIHIRIGSARDCEFYSKELSRIISDLEEPIIVYRCLHFLHRYY
Schizosaccharomyces_pombe  901  --------------------------------------------------------------------KYITWRLFAVSAHSNVCAARIYEHMGQAREAEFFYRQACSISEKMPFSC--FSAT-------

Arabidopsis_thaliana       1141 ---FSSA-----------------------------------------------------LGNLYHKKQCLDLAEKELQNAKEIL-----------------IANQRD----------------------
Oryza_sativa               1118 ---FASA-----------------------------------------------------LGQIYRKRQQWDTAEGELKYARDLL-----------------AQNATF----------------------
Chlamydomonas              1279 ---SCC-----------------------------------------------------CLADVACRRVDAAGAAASLQEAEEAVGRSCSPADASGASHGAIAAEAAL----------------------
Homo_Sapiens               1210 ---SCCTTQLFASSWGWQPPLIKSVPGSEPSKTQGQKRSGRGRQKLASAPLRLNNTSQKG---------------------LEGRGLPCTPKPPDRIRQAGP-HVPFT----------------------
Caenorhabditis_elegans     573  ---LEF---PMECNYSWMRPTIRK-----------------------------------------------------------------------PRVKATI-DCAVD----------------------
Saccharomyces_cerevisiae   878  MITEQTCLQNI------------------------------------------------TLGKANKA-FDYLDAEADITSLTMFL-----------------YDNKEFVKLEQSLVLYFGDQLEKTFLPN
Schizosaccharomyces_pombe  954  ---FQLR-----------------------------------------------------LCSLLTR-------AGKLEKGEKILFDLT-----------------------------------------

Arabidopsis_thaliana       1176 ---------FSCVKCKLKLEVT--LDKQLGD---------------------------------------------IS--RKQIDRVSQT--------------------DGFL-HAESLFSAALGKFCC
Oryza_sativa               1153 ---------ISCKLCKLTLDIS--LDVQAGD---------------------------------------------LFWSLYEKDFQKQS--------------------AGNLSNALGMYQSALDKLNG
Chlamydomonas              1331 ---------VCANLCAAQARRS--LLEGATDKAQAAATRGVGALTDVGGGAFGWLSVSAQSRLARLRAEVQLKLGQSVAASNITRDAVVHLQEA-----------------ASAAAADAAAESTEAGSAG
Homo_Sapiens               1293 ---------VFEEVCPTESKPE--VPQ-----------------------------------APRVQQR------------------------VQTRLKVNFSDDSDLEDPVSAE-AWLAEEPKRRGTAS
Caenorhabditis_elegans     603  ---------ILRAVSPFGRRPK--VEK-----------------------------------LE----------------------------------------------------KNL-----QPFDKE
Saccharomyces_cerevisiae   942  LWKLHLGKDIDDSICLSEYMPKNVINR-----------------------------------------------------------------------------------------VHNMWQKVMSQLEE
Schizosaccharomyces_pombe  980  ----------------------------------------------------------------------------------------------------------------------------------

Arabidopsis_thaliana       1228 SAWK-SCIRSHGEEIAEEIVIDRNGGEGLGHNSS-------------------------------------KT-----KLSIKEPPGNRGSRRGGRANKTCLS----KDQDLISEPTSRLTRSMR--HSL
Oryza_sativa               1208 TKLE-SPVDSYDKLKTTCIICSKYGKEPLAANDGVLPSCTVCANFSQ-------ASGDHSNE--FTALKFLKH-----KDSECCPPLDVKVKRTTR-NSSRLA----KEQNVEAHVKTRTRSSKRTAHMK
Chlamydomonas              1434 PEVL-WPVDY-----------------------GLLLSQQAAASRASGSGQP--GSGERGGA--PVVVGLCRH-----GAGQAGPG----------NND-------------------------------
Homo_Sapiens               1353 R-GR-GRARK-----------------------GLSLKTDAVVAPGSAPGNP--GLNGRSRRAKKVASRHCEERRPQRASDQARPG----------PEI-------------------------------
Caenorhabditis_elegans     631  R-F-------------------------------E-----------------------------------------------------------------------------------------------
Saccharomyces_cerevisiae   983  DPFFKGMFES-----------------------TLGIPSSLPVIPSTMPNNILKTPSKHST-----GLKLCDS---------------------PR-SSSMTP----RGKNIRQKFD-RIAAI-------
Schizosaccharomyces_pombe  980  -EAM-KSTDTYHKLLW------NYG------------AAEVCATKSE-------LDGAICHY--SECVKLLEI-----IKSEYYL-----------------FFNRNREKSLTKGIKRLSLSSQPTF--V

Arabidopsis_thaliana       1309 REQCQNRSNVPEVV------------------------SKKPNLCDRS-VGSRGERVLLD-TSNALPGFCICYKEKRQQCLSEEVTESGSLN----------NLVSLKWELCHRKLASSI----LVSLGK
Oryza_sativa               1318 GEKASTELHCKNGL------------------------SCSDNLSTDTLVRGKA-NCILDGVDQSIDYTCSIF--GCWNCLFVNTLNSGSIQ----------NILQFRWDCVWHHNHVSI----LLKIAK
Chlamydomonas              1489 ----------------------------------------------------------------------------------------------------------------------------------
Homo_Sapiens               1414 ------------------------------------------------------------------------------------------MR----------TIPEEELTDNWRKMSFEI----LRGSDG
Caenorhabditis_elegans     633  ----------------------------------------------------------------------------------------------------------------------------------
Saccharomyces_cerevisiae   1050 ----------------------------------------------------------------------------------------------------------------------------------
Schizosaccharomyces_pombe  1058 TESNTTEFD---DWSILQNTAANLLRLISMFELKRGNLEIAKALMTDS-------------------TKCSIA--SFFNIVSANILKSKLIVCEADSTLFGDPVLRTLPDSVISLPGISHKFQKNQSKTK

Arabidopsis_thaliana       1399 CLGDSGRIHLAHE-----ALLHSISVL-FK----STWSSHNQPSVSQLLEFIGKEVTRDVFAVDRAIILYNLCWLNLRNYHCRK-SRSICCDLFHIPF--T-KLVSWLMLAFVL--SGEV-PILFQ-KVS
Oryza_sativa               1407 ALGAHGGLHGAHK--IHNIYWQCISLLYFR----SLPQDCYRTYEHNLFGLIMDQSTGDFLISERAEILYSMSLFLLKGFLSEQ-SRDICCRFCSVQM--S-DVVPWLLKAFVL--SREN-PSLFQ-EVC
Chlamydomonas              1489 ---------TDMGSEAPDGKPKADQDQSSKRGGARGKAGAASSKASKATG--------PEG--------------------------NSATSTMSPGQASTSHVA-PLLLALRL--CWQV-PLAAT-HAA
Homo_Sapiens               1441 EDSASGGKTPAPGPEAASGEWELLRLDSSKK---KLPSPCPDKESDKDLG--------PRL--------------------------RLPSAPVATGLSTLDSICDSLSVAFRGI-SHCPPSGLYA-HLC
Caenorhabditis_elegans     633  ----------------------KV------------------------------------------------------------------------------------RLAMRNEMNHYG-HILYREWRC
Saccharomyces_cerevisiae   1050 -----------------------SKLKQMKE---LLESLKLDTLDNHELS--------KIS---------------------------SLSSLTLTILSNITSIHNAESSL-------------------
Schizosaccharomyces_pombe  1164 ALGENTGFRKGSK-----------RLDYLR------ERLKINL-QNV-----------------------RLSCEIIFSNAYERSSVC-----------------------------------VCR-EVN

Arabidopsis_thaliana       1511 R-LLASL-YLLSSSN------SE-FTFESDGNELSASHWVSFFHQASLG-THLSYHFISNLSQ----------------------KHKS-QCLSDKECTEATCSSCMVPEDLDLPRL-------------
Oryza_sativa               1523 R-LLACI-FLLATID----STAQ-LPLYSS-GSLSLNHWAAYFHQNSVG-TYLDCQYFAGLKS-----------------------------LLRKNDSKAALEDFSNASDESLSKF-------------
Chlamydomonas              1572 RQLLRLA-VALGLPH----AAAMFM-H-LANCASYPQ--------QQLLQEQMRRY-LAR-MTTAGAAAMAHDEAEAGSESDVDDERRGSGASASS---SGGCRTAAAAGDVREAALRLVA------ELC
Homo_Sapiens               1532 R-FLALC-LGHRDPY----ATAF-L-V-TESVSITCR--------HQLL-THLHRQ-LSK-A----------------------QKHRGSLEIADQ---LQGLSLQEMPGDVPLARIQRL----------
Caenorhabditis_elegans     657  R-LFAYVGRTSRDPW----EAAY-A-W-AESTQIGAR--------NAVQ-SRL----------------------------------------------------E------------KC----------
Saccharomyces_cerevisiae   1100 ---------------------------------------IT------------------------------------------------------------NFSLTDLPRHMPLLFDKVLNNIDNKNYRE
Schizosaccharomyces_pombe  1217 EL-ISYS-TIMQSALTTIGETTD-VD--SSSASFFLEI------PKALG-FHRRRE-AQKFRN----------------------QHKE-LHFSSL---EQI----------------------------

Arabidopsis_thaliana       1595 A--------------P---DR-----TQDL-----VQFAKEFFINLPSSTIICISLLGGALNQLLQELMHIRSPVCAWVLISRLNPESQPV-ATLLPVDSIVEDMSDDSANLSSTEATQVKSLKGPWLCP
Oryza_sativa               1602 FRF-----------SS---AD-----IGHL-----EIHIKEFFHKLPDVPIVCISMLEGDFVNVLGEILLLPSYFPAWMMLSRFDSTNKPI-TMLLPVDAISKETQHED-SCTKELDNLMRATDKNWQCP
Chlamydomonas              1676 DSGRGATCAGGDSTAA---LE-----RG-A-----EAWVREALGCLPADCVVASIME---------------DREDHVVLLGRLARGGPPL-LVMLPGPRESGTQG------------------------
Homo_Sapiens               1607 FSF-----------RA---LE-----SGHFPQPE-KESFQERLALIPSGVTVCVLALA----------TLQPGTVGNTLLLTRLEKDSPPV-SVQIPTGQNKL---------------------------
Caenorhabditis_elegans     696  KRG-----------LV---TM-----SG-------HDRFKTCVQSMPDEMTLVQIAMA----------------DDKTIYLVKLHADRDPI-IMPLAHYSQAV---------------------------
Saccharomyces_cerevisiae   1132 FRV-----------SSLIAPNNISTITESI---RVSAAQKDLMES---NLNINVIT-------------IDFCPITGNLLLSKLEPRRKRRTHLRLPLIRSNSRDL-DEVHLSF----------------
Schizosaccharomyces_pombe  1280 LNS------------R---LS-----IPDV-----RTFQDNFIDSLPSIWNVVSITI---------------NNSGEDLFISKIRKGHSPL-IFRLPLQRHNSRDADEE-ILVF----------------
```

```
Arabidopsis_thaliana       1697 WGTTVVDEVAPAFKSILEESHSSSS-T---TEEDTIESRGLWWKKRKKLNHRLGIFLRNLEASWLGPWRCLLLGEW------------------------------------------------------
Oryza_sativa               1706 WGYTIIDYVAPTFRKILEENFISLS-SATLTLNDGQANHVKWWSHRMKLNNHLDKMLKDMEESWLGPWKCLLLGYD------------------------------------------------------
Chlamydomonas              1752 GGDSHVGGCLGQLQRLLEDSGDSMRTD--QAAASSQNQKAQWWKARTALDASVSKLLQQLDSDCLGAWRCLLLPLPASHRGVLAGAAAAFVGEHVTGGGAAPAAGIATAVAQELLTVAMANMGQLGSNEV
Homo_Sapiens               1678 -----------HLRSVLNEFDAIQK-A--QKENSSCTDKREWWTGRLALDHRMEVLIASLEKSVLGCWKGLLLPSS------------------------------------------------------
Caenorhabditis_elegans     755  -------ELMDKFTFLLDEDEMIAK-------YPGDITPEEFWKRRKIVDGRMMTFVDEVQKHFLGVAASLLMPSG------------------------------------------------------
Saccharomyces_cerevisiae   1214 ------PEATKKLLSIINESNQTTS-VEVTNKIKTREERKSWWTTRYDLDKRMQQLLNNIENSWFNGVQGFFSPEV------------------------------------------------------
Schizosaccharomyces_pombe  1351 ------TKAQTELFRIISKSNQMAQ-N--GKHYTRREDKETWWKERRHLDQCLQQLLENIEISWLGGFKGIFNPHK------------------------------------------------------
```

```
Arabidopsis_thaliana       1768 ----------------------------------------------------SNYKLPDSA---------------------------------------------------------------------
Oryza_sativa               1780 ----------------------------------------------------LTDQHIEEA---------------------------------------------------------------------
Chlamydomonas              1880 GTVLRTVCGGISGGTSCNTDVAALAAELAAAQAAACMALLHLPQPAAGPTASPE-PAPAKPAPAPAAVAEPVEPVPVQPRARRPATAGASAATASSSARAVPGARARPGLAARPVATEAVRRTTAAAAAN
Homo_Sapiens               1740 ----------------------------------------------------EE-PGPAQE---------------------------------------------------------------------
Caenorhabditis_elegans     817  ----------------------------------------------------QLGPKAAEL---------------------------------------------------------------------
Saccharomyces_cerevisiae   1283 ----------------------------------------------------VDNSLFEKFK--------------------------------------------------------------------
Schizosaccharomyces_pombe  1418 ----------------------------------------------------IDTSLFAKFS--------------------------------------------------------------------
```

```
Arabidopsis_thaliana       1777 -------------------------------QKKLVND----LKSKCK-MEVNEMLLKVILGGGT-DNFKGEACVAQLSLRNGCYVGRGG-------YLYEEDSCKTPTAASN--------ISESRHELA
Oryza_sativa               1789 -------------------------------LTNLIAG----LESEFK-FEVNPVLIKVILGGAM-SVDEVQDCVSQLISYKG-YFGRGG-------CC-GKDRLRA--LSSC--C-----IESEALETV
Chlamydomonas              2009 IAASLEALTVEDDAASTSAAPAADQPARSRRAPRLAAMQSESAASAASKAPPTRGL--VAASKA-------------------------G-------ARTARA-APA--PAATLATTERV-ATLSKARTP
Homo_Sapiens               1748 -------------------------------ASRLQEL----LQDCGW-KYPDRTLLKIMLSGA-------------------------G-------ALTPQD-IQA--LAYGLCP-----TQPERAQ--
Caenorhabditis_elegans     826  -------------------------------AIKIHK-----LSKGGL-LLGEA---KEMVYQS-------------------------K-------LMDAKS-WEA--LILRFCE--MR-TTDEKFKSF
Saccharomyces_cerevisiae   1293 --------------------------------DKFYEI----LHQN---------LPSRKLYGNPAMFIKVEDWVIELFLK----L-NPQEIDFLSKMEDLIY-FVL--DILLFHG--EENAYDEIDFSM
Schizosaccharomyces_pombe  1428 --------------------------------SQFQNI----IAKN---------FNMDKKTPVPTL----SPEILELFIT----LGKPGYEGYEQLLEDLIY-FIL--DIFQFRG--LHFAYDEIDTDQ
```

```
											*  esp1-P1946L
Arabidopsis_thaliana       1856 LKLI-H-DAA--SKLG----QQDGHENREPIILVLDPEVQMLPWENIPILRKQE-VYRMPSVGCISAVL-KKR-SLQGEPAKSHVASFPLIDPLDSFYLLNPGGDLTDTQVTFESWFR-----DQNFEGK
Oryza_sativa               1865 ECLI-K------STVN----ELIEPVDRDPVIFVLDTNVQMLPWENLPALRNQE-IYRMPSIGSVFLAL-TRS-NNYWKDARVIAPPFPVIDPFNAFYLLNPSGDLSSTQEEFDQMFK-----NYEWKGK
Chlamydomonas              2101 APAM-A-S--EVAQQGTVATSNGGAAGSCPLLLVLSPCLHALPWESMPCLRGRS-VSRVLSLPACCGAA-AAAFTSGG------GRSSPAALPTSAFYLLNPSGDLADTQSAFQQLLEA----QAGWQGV
Homo_Sapiens               1800 -ELL-N-EAV--GRLQ-----GLTVPSNSHLVLVLDKDLQKLPWESMPSLQALP-VTRLPSFRFLLSYSIIKE-YGAS------PVLSQGVDPRSTFYVLNPHNNLSSTEEQFRANFSS----EAGWRGV
Caenorhabditis_elegans     879  LPLM-HRNSVEVMNQD-----DSIVTEKKYTYLVICPHLSQFCWERLPIFDEYPYVGRQVSIHSTFSQL-EAM-KSQE------KQIPLQIDVQNAYYILDPDNNLGETQKRMVEYIN-----KFNWEGT
Saccharomyces_cerevisiae   1369 LHVQLE-EQI--KKYR----ATMTTNSIFHTFLVVSSSCHLFPWECLSFLKDLS-ITRVPSYVCLNKLL-SRF-HYQL--------PLQVTIEDNISMILNPNGDLSRTESKFKGMFQKIIDAKPSSQLV
Schizosaccharomyces_pombe  1501 LSMDLQ-DALN-AYFN----NYVSEENRSHTVLVLDKSVHQFPWESLPCLNRQS-VSRVPSLSILRDIL-SQS-FVVN------G-EYVEVRKEAGSYILNPSLDLKHTQEMFEHKLV-----EGGWKGL
```

```
									  *  esp1- A2047T
Arabidopsis_thaliana       1970 AGSEPSAIELTEALETHDLFLYFGHGSGAQYIPRREIEKLDNCSATFLMGCSSGSL-WLKGCYIPQGVPLSYLLGGSPAIVATLWDVTDRDIDRFGKALLEAWLQER-SDSSSEGGCSQCESLANDLAAM
Oryza_sativa               1976 AGYAPTAEELVLALRNHDLFLYFGHGSGTQYVSGKEIEKLDNCAAALLMGCSSGTL-RCKGCYAPQGAPLSYLSAGSPAVIANLWDVSDKDIDRFSKALLGSWLQEN-FVA--AKNCSKCCQLTREFESM
Chlamydomonas              2215 VGQQPSARALLAALANHEVFLFLGHGSGEQYLPLPALRKLQRCAAAVLMGCSSGRL-RLHGAYDPAGAVVAYAVAGSPAVVANLWDVTDRDIDRYCQALLRNWLGCADPQA--AAAAAAAGQEDEEQEAQ
Homo_Sapiens               1909 VGEVPRPEQVQEALTKHDLYIYAGHGAGARFLDGQAVLRLSCRAVALLFGCSSAAL-AVRGNLEGAGIVLKYIMAGCPLFLGNLWDVTDRDIDRYTEALLQGWLGAG-----------------------
Caenorhabditis_elegans     990  VGSAPKSNEISAALSQRDAFFFIGHGSGSSVMPRSVLKQSTCNAISLLMGCGSVRTIPQALGFDGKTAILDYAMAKCPLIVGCLWTVTDGEIDRFLIRMIDDCFEDS--KS--LTGID------------
Saccharomyces_cerevisiae   1481 MNEKPEEETLLKMLQNSNLFVYIGHGGGEQYVRSKEIKKCTKIAPSFLLGCSSAAM-KYYGKLEPTGTIYTYLLGGCPMVLGNLWDVTDKDIDKFSEELFEKMGFRCNTDD--LNG--------------
Schizosaccharomyces_pombe  1610 IASQPSNRDFIKMLSGNDFFLYFGHGGGEQYTTSYDLATLKRCAVTILMGCSSGAL-YECGSFEPWGTPLDYLSAGCPTLVANLWDVTDKDIDRFSLKMLESWGLFENKAP--FV---------------
```

```
							        *  esp1- P2156S
Arabidopsis_thaliana       2098 TL--KGTKRSRKPSSRNKPAQSDVDGSGKIECNHKHRRKIGSFIAAARDACNLQYLIGAAPVCYGVPTGI---------------------------------------T--------------------
Oryza_sativa               2102 TIAVEGNGRPRRRGTRGKKSERMNNCSKRC---TCGNRRVASYLSEARRACRLPLMIGGSPVCYGVPTII---------------------------------------R--------------------
Chlamydomonas              2342 PVGW---------------------------------AGLGQAVVSSRGACRLPHLIGAAPVCYGLPLHQ------------------------------------------------------------
Homo_Sapiens               2014 --PG---------------------------------APLLYYVNQARQAPRLKYLIGAAPIAYGLPVS---------------------------------------LR--------------------
Caenorhabditis_elegans     1103 --KL---------------------------------RQLSEAMHEARSKARLKYLTGAAVVMYGLPVVAKQTTPFVEKDQRNLPQTPKTSARTSMRMETVPKTPKQEFVTSKSVPMTPIFSNNENKSPS
Saccharomyces_cerevisiae   1593 --NS---------------------------------LSVSYAVSKSRGVCHLRYLNGAAPVIYGLPIKF---------------------------------------VS-------------------
Schizosaccharomyces_pombe  1721 --NS---------------------------------TSICTAVSESRSCCHLRYLNGAAPVIYGIPAYI---------------------------------------IP-------------------
```

```
Arabidopsis_thaliana       2166 ------------------------------------------------RKKGIDALLPSSSR--
Oryza_sativa               2169 ------------------------------------------------KK--------------
Chlamydomonas              2378 ----------------------------------------------------------------
Homo_Sapiens               2050 ----------------------------------------------------------------
Caenorhabditis_elegans     1199 RARMPSRVLKTPRQVKTFQEEDDEAPKRSTTRQLKPLVAPPIPATPTTRTTRSSARTPSRSRNL
Saccharomyces_cerevisiae   1630 ----------------------------------------------------------------
Schizosaccharomyces_pombe  1758 ----------------------------------------------------------------
```

**Figure S3. Alignment of separase proteins from distant eukaryotes.**

The separase protein from *Arabidopsis thaliana* (AAW32909.1), *Oryza sativa* (XP_015626143.1), *Chlamydomonas* *reinhardtii* (PNW88431.1), *Homo Sapiens* (Q14674.3), *Caenorhabditis elegans* (AAK77200.1), *Saccharomyces cerevisiae* (NP_011612.3) and *Schizosaccharomyces pombe* (CAA21959.1) are shown aligned. Amino acids similar in at least three proteins are shaded in colour. The position of the four missense mutations identified in this study are indicated by red stars. The SD (substrate-binding domain) and CD (catalytic domain) are underlined in green and turquoise, respectively (Luo and Tong, 1017). T-COFFEE (<https://toolkit.tuebingen.mpg.de/#/tools/tcoffee>) and Bioedit 7.2.5 were used to produce the alignment.

.
